# Distinct DNA-binding surfaces in the ATPase and linker domains of MutLγ determine its substrate specificities and exert separable functions in meiotic recombination and mismatch repair

**DOI:** 10.1101/108647

**Authors:** Corentin Claeys Bouuaert, Scott Keeney

## Abstract

Mlh1-Mlh3 (MutLγ) is a mismatch repair factor with a central role in formation of meiotic crossovers, presumably through resolution of double Holliday junctions. MutLγ has DNA binding, nuclease, and ATPase activities, but how these relate to one another and to *in vivo* functions are unclear. Here, we combine biochemical and genetic analyses to characterize *Saccharomyces cerevisiae* MutLγ. Limited proteolysis and atomic force microscopy showed that purified recombinant MutLγ undergoes ATP-driven conformational changes. *In vitro*, MutLγ displayed separable DNA-binding activities toward Holliday junctions (HJ) and, surprisingly, single-stranded DNA (ssDNA), which was not predicted from current models. MutLγ bound DNA cooperatively, could bind multiple substrates simultaneously, and formed higher-order complexes. FeBABE hydroxyl radical footprinting indicated that the DNA-binding interfaces of MutLγ for ssDNA and HJ substrates only partially overlap. Most contacts with HJ substrates were located in the linker regions of MutLγ, whereas ssDNA contacts mapped within linker regions as well as the N-terminal ATPase domains. Using yeast genetic assays for mismatch repair and meiotic recombination, we found that mutations within different DNA-binding surfaces exert separable effects *in vivo*. For example, mutations within the Mlh1 linker conferred little or no meiotic phenotype but led to mismatch repair deficiency. Interestingly, mutations in the N-terminal domain of Mlh1 caused a stronger meiotic defect than *mlh1Δ*, suggesting that the mutant proteins retain an activity that interferes with alternative recombination pathways. Furthermore, *mlh3Δ* caused more chromosome missegregation than *mlh1Δ*, whereas *mlh1Δ* but not *mlh3Δ* partially alleviated meiotic defects of *msh5Δ* mutants. These findings illustrate functional differences between Mlh1 and Mlh3 during meiosis and suggest that their absence impinges on chromosome segregation not only via reduced formation of crossovers. Taken together, our results offer insights into the structure-function relationships of the MutLγ complex and reveal unanticipated genetic relationships between components of the meiotic recombination machinery.

**Author summary:** Sexual reproduction involves the fusion of two gametes that each contain half of the DNA from each parent. These gametes are generated through a specialized cellular division called meiosis. During meiosis, the cell faces the challenge of identifying the appropriate pairs of chromosomes that need to be separated. This involves an elaborate mechanism whereby the parental chromosomes recombine and form crossovers, i.e. exchange DNA fragments. These crossovers are thus important for the accurate segregation of chromosomes and are also fundamental to evolution because they help shuffle linkage groups from one generation to another. Here, we have studied a complex of proteins called MutLγ that is important for the formation of crossovers, and is also involved in an unrelated mechanism that repairs mistakes that spontaneous arise in DNA when it is synthesized. We uncovered intriguing features of the interaction of this complex with DNA. In addition, by studying a collection of mutants of MutLγ, we identified mutants that affect one biological function but not another. For example, surprisingly, we found mutations that decrease the frequency of crossovers but did not affect chromosome segregation as much as expected. Taken together, our findings allow us to reconsider the ways in which we think about these processes.

## Introduction

During meiosis, cells undergo DNA recombination to form crossovers between homologous pairs of chromosomes (homologs). Crossovers promote accurate segregation of homologs at the first meiotic division and increase genetic diversity by breaking up linkage groups [1].

Recombination is initiated by DNA double-strand breaks made by Spo11 [2–4], which remains covalently attached to the DNA and is released by endonucleolytic cleavage [5]. Double-strand breaks are then resected to form 3′ single-stranded tails, which serve as a substrate for strand exchange proteins to invade a homologous template [6, 7]. Subsets of these initial invasions further mature, after DNA synthesis and capture of the second end, into double Holliday junction (dHJ) intermediates, which are finally resolved into crossovers [1, 8, 9]. Because crossovers are crucial to meiosis, the cell tightly controls their number and distribution [10–13].

MutLγ is important for crossover formation in most organisms, including yeast and mammals [14–18]. MutLγ is believed to catalyze the nuclease reaction that resolves the dHJ intermediate into a crossover [17, 19]. Other proteins implicated in regulated crossing over include the ZMMs (Zip2-Zip3-Zip4-Spo16, Msh4-Msh5, Mer3), a biochemically and functionally diverse group of proteins that channel recombination intermediates toward a crossover fate [20, 21]. In addition to the major MutLγ- and ZMM-dependent pathway, another crossover pathway in *S. cerevisiae* depends on the structure-specific nuclease Mus81-Mms4 [22, 23]. Mus81-Mms4 is thought to be responsible for ~15% of crossovers in wild-type yeast but can partially substitute when MutLγ is compromised [17, 22]. Several additional systems can also take apart dHJ intermediates, namely the structure-specific nucleases Yen1 and Slx1/Slx4 and the combined activity of Sgs1 helicase plus Top3 topoisomerase in complex with Rmi1 protein [24–27]. These latter systems are largely cryptic in wild-type cells and presumably contribute failsafe mechanisms that can scavenge recombination intermediates that escape the normal resolution pathways [17, 28, 29].

Mlh1 and Mlh3 are also involved in post-replication mismatch repair (MMR) [30]. Mlh1 and Pms1 form the central MLH complex in yeast (MutLα) that is targeted to DNA mismatches by an MSH complex (Msh2-Msh6 or Msh2-Msh3) and that introduces DNA nicks to initiate degradation and repair of a mismatch-containing strand [31, 32]. Mlh3 also participates in MMR along with Mlh1, but in a minor role [33, 34]. MutLα also functions in repair of mismatches formed within the heteroduplex DNA intermediates of meiotic recombination [14, 35]. Furthermore, Mlh1 along with Mlh2 forms a third heterodimeric complex, MutL_β_, which has as yet poorly understood functions in controlling meiotic gene conversion patterns [35, 36]. Importantly, however, MutLγ is the only MLH complex critical for meiotic crossing over *per se*, as MutLα and MutL_β_ are fully dispensable for formation of normal crossover numbers [14, 35, 36].

The ZMM proteins Msh4 and Msh5 (MutSγ) are also related to MMR factors but play no role in MMR [37, 38]. By analogy with MMR, it has been proposed that MutSγ binds to recombination intermediates and recruits MutLγ to catalyze crossover formation [17]. Experimental evidence consistent with this idea includes cytological studies in mice and other organisms that show that the appearance of MutSγ precedes the appearance of MutLγ foci and that the number, timing and distribution of MutLγ foci correlate with chiasmata, indicating that MutLγ marks crossover sites [39–43]. *In vitro*, human MutSγ binds to Holliday junctions (HJ) and related branched structures [44]. However, MutLγ alone also binds specifically to HJs, independently of MutSγ [45]. In addition, nuclease activity of MutLγ has been demonstrated using plasmid DNA substrates, but HJ resolution activity has not yet been reconstituted [45, 46]. Thus, key steps in meiotic crossing over remain poorly understood.

In addition to DNA binding and cleavage activities, MutLγ possesses ATP binding and hydrolysis activities that appear to be essential for its MMR and meiotic functions, although controversy remains as to whether ATP hydrolysis is required in meiosis [18, 46–49]. As in other MLH proteins, MutLγ ATPase activity is carried within an N-terminal GHKL domain, which is connected to the C-terminal domain by a flexible linker [50]. The nuclease active site is in the C-terminal domain of Mlh3, which dimerizes with the Mlh1 C-terminal domain [19, 45, 46]. In bacterial MutL and eukaryotic MutLα, cycles of ATP binding, hydrolysis, and nucleotide release modulate the conformational state of the complexes through dimerization of the N-terminal domains. These structural changes are proposed to act as a molecular switch to transduce signals between mismatch recognition factors and repair [51–53]. Nevertheless, how the DNA-binding, nuclease, and ATPase activities of eukaryotic MLH complexes relate to one another and to their MMR and/or crossover-promoting properties are unclear. We set out in this study to address this lack by combining genetic approaches with detailed biochemical characterization of the DNA-binding properties of *S. cerevisiae* MutLγ.

## Results

### Purification of MutLγ, catalytic activities and ATP-mediated conformational changes

To study the biochemical properties of *S. cerevisiae* MutLγ, we purified N-terminally-tagged Mlh1-Mlh3 heterodimers from baculovirus-infected insect cells (Materials and Methods) (**Fig 1A**). We verified that the tagged proteins are functional in yeast, using strains that express identically tagged versions of Mlh1 (^*HisFlag*^*mlh1*) and Mlh3 (^*HisFlag*^*mlh3*) from their endogenous loci. Using genetic assays (described below), we found that the ^*HisFlag*^*mlh1* strain displayed a possible mild increase in chromosome missegregation that was not statistically significant (0.6%, p = 0.057, Fisher’s exact test (two tailed *P* value)) and retained wild-type levels of crossing over and MMR (**Fig 1B, C**). The ^*HisFlag*^*mlh3* strain had a crossover defect (80% of wild-type levels, p < 0.05, G test) but essentially wild-type levels of chromosome missegregation (0.3%, p = 0.29 Fisher’s exact test) and wild-type levels of MMR (**Fig 1B, C**). The biochemical experiments presented below used MutLγ complexes tagged on Mlh1, unless stated otherwise.

**Fig 1:**
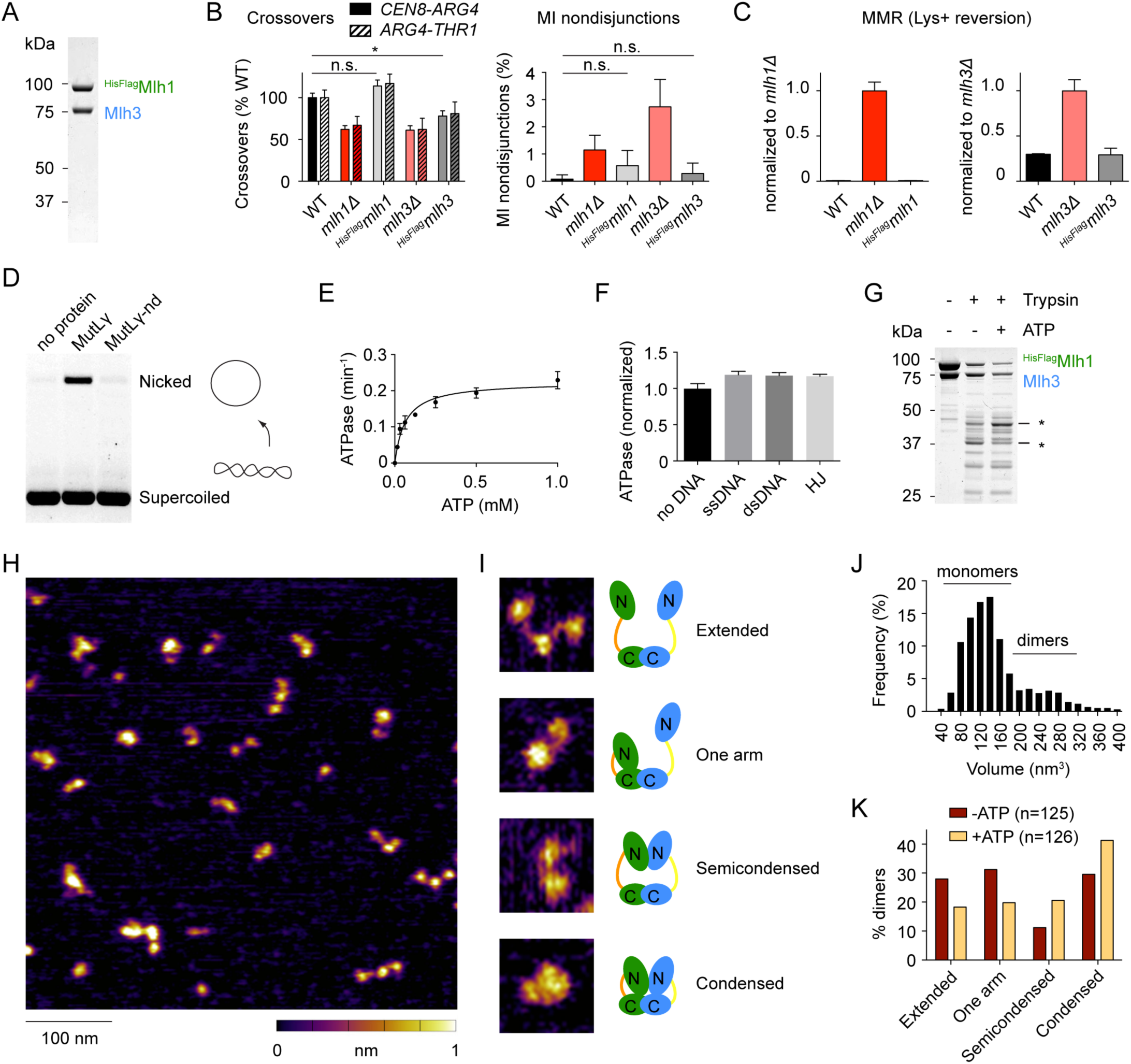
Purification and *in vitro* characterization of the *S. cerevisiae* MutLγcomplex. A. Purification of affinity-tagged *S. cerevisiae* Mlh1-Mlh3 heterodimers from baculovirus-infected insect cells. A Coomassie-stained SDS-PAGE gel is shown; 2 g of protein was loaded. B. Effects on meiotic recombination of N-terminal tagging of Mlh1 (*HisFlagmlh1*) and Mlh3 (*HisFlagmlh3*). Crossover frequencies in two test intervals (*CEN8-ARG4* and *ARG4-THR1*) and MI nondisjunctions frequencies were measured by the spore-autonomous fluorescence assay (described in detail in Fig 4 below; values for *mlh1*and *mlh3*are reproduced from Fig 4 for comparison). Error bars represent standard errors for crossovers and 95% confidence interval of the proportion for MI nondisjunction. Statistically significant differences are indicated with asterisks (*, p<0.05). Selected non-significant differences are highlighted (n.s.). C. Effects on MMR of N-terminal tagging of Mlh1 and Mlh3. Graphs show Lys+ reversion frequencies normalized to the wild-type value. See text associated with Fig 5 for details on the assay. Error bars are standard deviation of the mean (n = 3–6). Values for *mlh1*and *mlh3*are reproduced from Fig 5 for comparison. D. DNA cleavage activity of MutL. Reactions containing 100 nM MutLγand 5.7 nM pUC19 plasmid were incubated at 30°C for 1 hour, then stopped, separated on an agarose gel, and stained with ethidium bromide. The nuclease-dead mutant (MutL-nd) carries the active site mutation D523N on Mlh3. E. Michaelis-Menten data of MutLγATPase activity (Vmax = 0.225 0.012 min-1; Km = 0.064 0.013 mM; mean ± SE). Error bars show the range from two independent experiments. F. Effect of DNA on the ATPase activity of MutL. Substrates are identical to the ones used for binding assays in Fig 2C (see Materials and Methods). Reactions contained 280 nM MutLγwith 4.8 pg DNA (2, 1 and 0.5 M ssDNA, dsDNA and HJ, respectively). Error bars show the standard deviations from three independent determinations. Data were normalized to the mean of the ATPase activity in the absence of DNA (kcat = 0.112 0.005 min-1). G. Partial proteolysis of MutL. A Coomassie-stained SDS-polyacrylamide gel is shown. Asterisks indicate some of the protein fragments that are different in the presence or absence of ATP (5 mM). H. AFM imaging of MutL. The image illustrates the heterogeneity of the size and configurations of MutLγparticles observed I. Examples of the four configurations of dimeric MutLγparticles observed by AFM. J. Volume analysis of MutLγparticles. The predicted volume of the heterodimer (175 kDa) is 257 nm3 according to the equation: Vc = (M0/N0)(V1+*d*V2), where M0 is the molecular weight, N0 is Avogadro’s number, V1 and V2 are specific volumes for protein (0.74 cm3 g-1) and water (1 cm3 g-1) and *d* is the extent of protein hydration (0.4 g H2O/g protein) [75]. K. Classification of MutLγparticles in the absence and presence of 1 mM ATP. A cutoff of 170 nm3 was chosen to classify particles as dimers (see panel J). The difference between distributions is statistically significant by G test, p = 0.0082.

Purified MutLγ displayed the expected nuclease activity on a supercoiled plasmid substrate. Under these conditions, wild-type MutLγ converted ~15% of the supercoiled substrate to nicked product, but nicking was undetectable when Mlh3 carried the nuclease domain mutation D523N (MutLγ-nd) (**Fig 1D**). This is in agreement with published results [45, 46]. To assay for ATPase activity, MutLγ was incubated with [α-^32^P]-ATP, then ATP hydrolysis products were separated by thin layer chromatography. MutLγ exhibited a low ATPase activity (k_cat_ = 0.1 min^-1^; **Fig 1E**), similar to other MLH complexes [46, 51, 54]. ATPase activity was not significantly stimulated by DNA (**Fig 1F**), as reported previously [46].

To address whether MutLγ undergoes ATP-driven conformational changes, we performed partial trypsin digestions of MutLγ in the presence and absence of ATP (**Fig 1G**). Gel electrophoresis of proteolysis reactions revealed that the presence of ATP results in specific changes in the pattern of trypsin cleavage fragments (see asterisks in **Fig 1G**).

We imaged the protein particles by atomic force microscopy (AFM) to gain insights into the molecular organization of the complex (**Fig 1H**). A volume analysis of the particles revealed that Mlh1 and Mlh3 exist as an equilibrium between monomers and dimers, with about one third of dimers at the concentration of this experiment (10 nM) (**Fig 1J**). Dimers exhibited different configurations that could be classified as extended, one-arm folded, semi-condensed, and condensed (**Fig 1I**), as previously reported for yeast and human MutLα [53]. In the presence of 1 mM ATP, the population of semi-condensed and condensed particles increased at the expense of extended and one-arm folded particles (**Fig 1K**). This is consistent with the idea of a molecular switch modulated by ATPase cycles [51, 53].

### MutLγ has distinct binding specificities for ssDNA and Holliday junction substrates

We assembled protein-DNA complexes of MutLγ using DNA substrates immobilized on streptavidin-coated beads (**Fig 2A**). MutLγ was efficiently pulled-down on beads coated with ssDNA (80 nt) or HJs (40-bp arms), but not double-stranded DNA (dsDNA, 80 bp), as revealed by SDS-PAGE of the bound fractions detected by silver staining and anti-Flag western blotting (**Fig 2A**).

**Fig 2:**
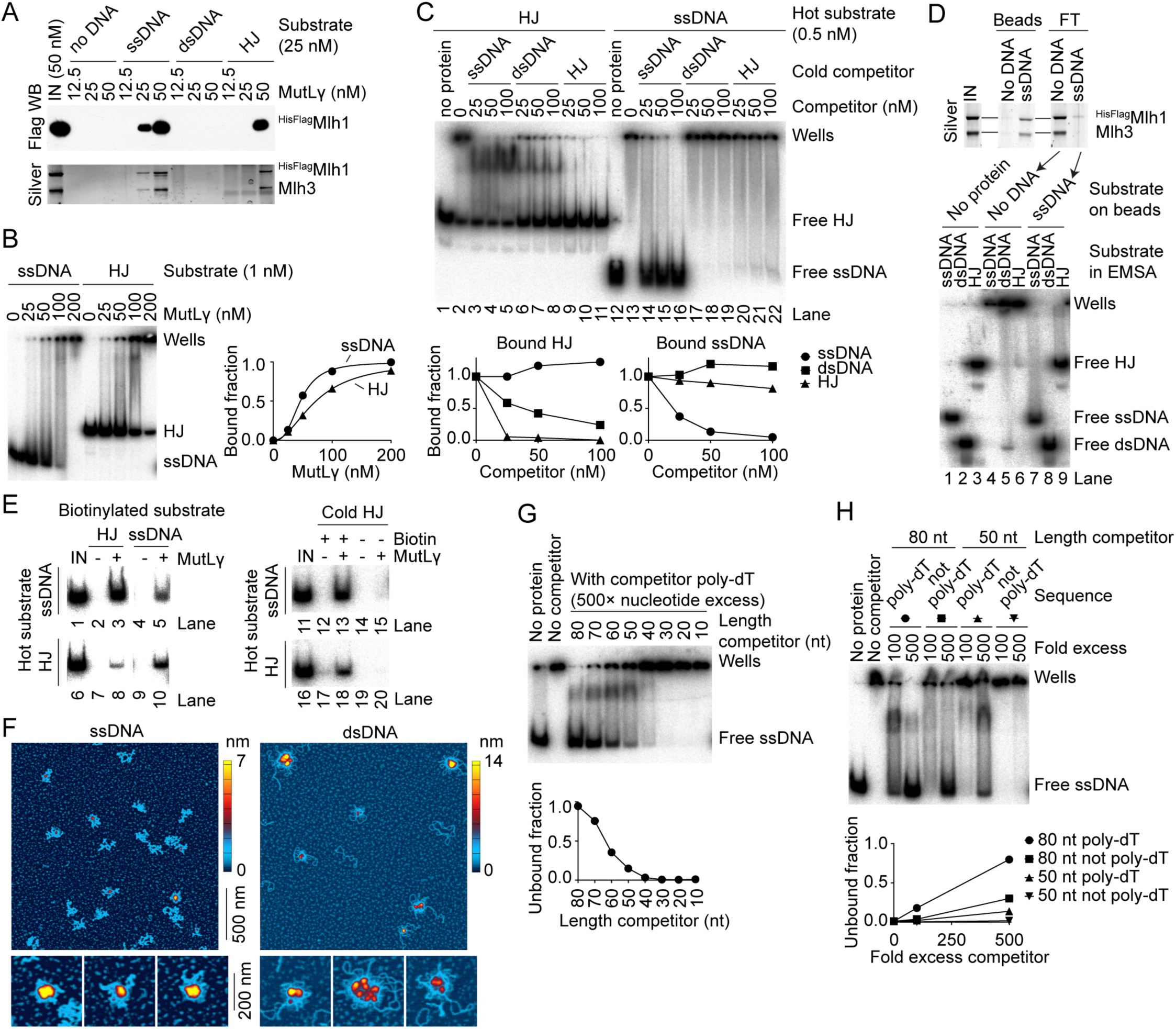
DNA-binding properties of MutL. A. Binding of HisFlagMlh1-Mlh3 to immobilized DNA substrates. Pulled-down proteins were separated by SDS-PAGE and detected by anti-Flag western blotting (top) and silver staining (bottom). Substrates are 80 nt poly-dT (ssDNA), 80 bp dsDNA, or HJ containing 40 bp arms. B. Binding of MutLγto 5′ end-labeled ssDNA (left) and HJ (right). MutLγwas titrated in the presence of 1 nM substrate. An autoradiograph of a 5% TAE-polyacrylamide gel is shown. Quantification of unbound substrates is plotted. The quantifications fit Hill equations with slopes of 2.9 ± 0.27 for ssDNA and 2.0 ± 0.05 for HJ, respectively and KD of 45.9 1.6 nM for ssDNA and 72.9 ± 1.3 nM for HJ, respectively (mean ± SE). C. MutLγhas specific and separable DNA-binding activities for ssDNA and HJs. Binding reactions contained 100 nM MutL, 0.5 nM end-labeled HJ (left) or ssDNA (right) and the indicated amount of cold competitor. An autoradiograph of a 5% TAE-polyacrylamide gel is shown. Quantification of the bound fractions is plotted. The competitor titrations were repeated ≥3 times with similar results. D. MutLγssDNA and HJ-binding activities are present in the same pool of proteins. The ssDNA-binding activity of MutLγwas depleted by pull-down on ssDNA-coated streptavidin beads (top) and the flow-through (FT) was tested in an EMSA for presence of ssDNA, dsDNA and HJ-binding activity (bottom). E. Simultaneous binding to multiple substrates. MutLγ(50 nM) was incubated with equimolar concentrations of biotin-labeled and radio-labeled substrates (25 nM each). Nucleoprotein complexes were pulled down on streptavidin beads and the presence of radiolabeled DNA was assayed by gel electrophoresis and autoradiography. The control experiment in the right panel compares pulldown efficiency with and without biotin label on the cold HJ substrate. F. MutLγforms higher-order complexes. AFM images of MutLγbound to ssDNA (10.4 103 nt, left) or supercoiled DNA (2.7 kb, right). Images are 2 m 2 m. Zoomed images illustrate the structural diversity of the higher-order complexes. G. MutLγhas a preference for long ssDNA substrates. Binding of MutLγ(100 nM) to radiolabeled ssDNA (poly-dT) was competed with a 500excess (in nt) of ssDNA substrates ranging from 10 to 80 nt. Quantification of the unbound fraction is plotted. H. The ssDNA-binding activity of MutLγdepends on the size and sequence of the substrate. Binding to an 80 nt radiolabeled poly-dT substrate was competed with 80 nt or 50 nt substrates that are either poly-dT or a different sequence. The 50 nt non-poly-dT substrate is identical to, the ssDNA previously reported not to be bound by MutLγ[45].

To measure the affinity of MutLγ for ssDNA and HJ substrates, we titrated MutLγ in the presence of 1 nM 5′ end-labeled ssDNA or HJ substrates and separated the protein-DNA complexes by gel electrophoresis (**Fig 2B**). Substrate binding reached over 90% completion at ~100–200 nM MutLγ. The affinity of MutLγ for the ssDNA substrate was higher than for the HJ substrate, with a K_D_ of 46 nM and 73 nM, respectively. The titrations fit Hill equations with slopes of 2.9 and 2.0 for ssDNA and HJ substrate, respectively, indicating that MutLγ binds cooperatively to both DNA substrates (**Fig 2B**, right). In addition, most of the nucleoprotein complexes remained stuck in the wells even when unbound substrate remained, consistent with cooperative formation of higher-order multimers.

To address the specificity of MutLγ for the ssDNA and HJ substrates, we performed competition experiments in an electrophoretic-mobility shift assay (EMSA) (**Fig 2C**). Nucleoprotein complexes were assembled with 100 nM MutLγ, 0.5 nM 5′ end-labeled HJ or ssDNA substrate, and 0–200-fold molar excess of cold competitor. In the absence of competitor DNA, MutLγ bound most of the DNA substrates and generated well-shift complexes (**Fig 2C**, lanes 2 and 13). With a 5′ end-labeled HJ substrate, binding was efficiently competed with a cold HJ competitor (lanes 9-11). In contrast, dsDNA and ssDNA competed less efficiently (lanes 3-8), indicating that the binding activity is specific to the HJ substrate. Conversely, with a 5′ end-labeled ssDNA substrate, binding was efficiently competed with cold ssDNA (lanes 14-16), but dsDNA and HJ competed less efficiently (lanes 17-22), indicating that binding to the ssDNA substrate is also specific. This does not contradict earlier reports that observed MutLγ binding on dsDNA [45, 46]. It indicates instead that, compared to dsDNA, MutLγ binds specifically to both ssDNA and HJ substrates.

To further investigate these two distinct DNA-binding activities, we depleted the protein sample of the ssDNA-binding activity by preincubating MutLγ with ssDNA-coated streptavidin beads and tested the supernatant for DNA binding (**Fig 2D**). We found that the HJ-binding activity of MutLγ had also been depleted (compare lanes 6 and 9), which suggests that both activities are present in the same pool of MutLγ molecules rather than representing distinct subpopulations of the protein.

If so, a further implication is that MutLγ can bind multiple substrates simultaneously. To address this, we incubated 50 nM MutLγ in the presence of 25 nM of a radiolabeled substrate and 25 nM of a biotin-labeled substrate. After complex assembly, biotin-labeled DNA was pulled down with streptavidin-coated beads. The beads were washed extensively, then deproteinized, separated by gel electrophoresis, and radiolabeled DNA was detected by autoradiography (**Fig 2E,** left). In the presence of MutLγ, radiolabeled substrates were pulled down with streptavidin beads, indicating that biotin-labeled and radio-labeled substrates were bound simultaneously by MutLγ (lanes 3, 5, 8 and 10). Pulldown of radiolabeled substrate was dependent on the presence of the biotin label, thus cannot be ascribed to nonspecific interactions between MutLγ and the streptavidin beads (**Fig 2E,** right, compare lanes 13 and 18 to lanes 15 and 20). These findings are consistent with individual MutLγ heterodimers binding more than one substrate simultaneously, or binding of multiple substrates through multiple proteins bound to one another.

To further investigate formation of higher-order nucleoprotein complexes, we used AFM to image MutLγ particles bound to ssDNA or plasmid substrates (**Fig 2F**). Reactions were assembled with 2 nM DNA and 10–40 nM MutLγ, then plated on mica slides and dried. Large protein-DNA structures were visible with both substrates (compare scales in **Fig 2F** with protein alone images in **Fig 1H**). These higher order structures were similar to those previously reported with MutLα [55].

A preference of MutLα for large DNA substrates has been observed and suggested to take part in the cooperative assembly of higher-order nucleoprotein complexes [55]. To test whether this was also the case for MutLγ, we performed substrate competition reactions where binding of MutLγ to an 80 nt labeled ssDNA substrate was competed with 500-fold excess (in nucleotides) of cold ssDNA substrates ranging from 10 to 80 nt (**Fig 2G**). No competition was observed with substrates smaller than 30–40 nt. Competition increased with larger substrate size and reached ~50% with 60–70 nt substrates.

A previous study also reported binding of MutLγ to HJs but did not detect binding to ssDNA [45]. Two factors are responsible for this apparent discrepancy: the size and the sequence of the ssDNA substrate used. When 80 nt or 50 nt substrates that were either poly-dT or another (non-poly-dT) sequence were compared as competitors in an EMSA with radiolabeled ssDNA substrate, the best competitor was the longest poly-dT substrate, whereas the shorter non-poly-dT sequence was almost completely ineffective under these conditions (**Fig 2H**). It is possible that, in addition to an affinity for larger ssDNA substrates, MutLγ prefers ssDNA that is less able to form secondary structures.

### Mapping the protein-DNA binding interfaces of MutLγ by hydroxyl radical footprinting

We mapped the ssDNA and HJ-binding interfaces of MutLγ by FeBABE footprinting (**Fig 3**). The technique takes advantage of an Fe^3+^ ion chelated by an EDTA moiety that can be chemically conjugated to a sulfhydryl group present on phosphorothioate-modified DNA substrates [56]. Upon activation with hydrogen peroxide, FeBABE generates hydroxyl radicals that cleave peptide or DNA chains within ~15–20 Å of the FeBABE binding site. Using terminally tagged proteins, the sizes of the protein fragments can be estimated by western blotting and provide an estimate of positions in the protein that were in proximity to the DNA.

**Fig 3:**
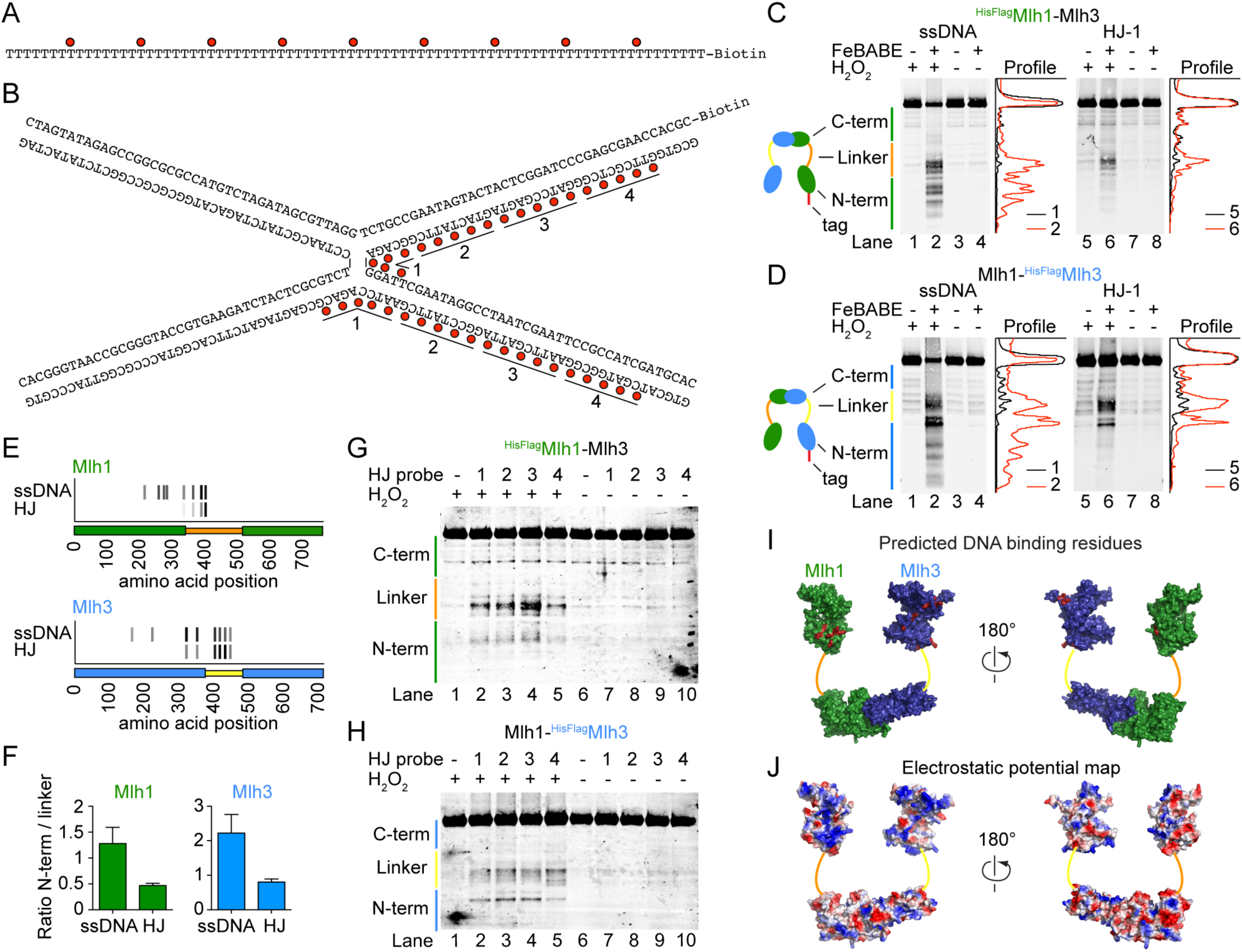
Mapping the DNA-binding interfaces of MutLγ by hydroxyl radical footprinting. A, B. Sequences of ssDNA (A) and HJ (B) substrates highlighting the positions of the FeBABE moieties (red dots). HJ substrates were labeled on two strands within the numbered segments: labeling in segments 1 was used for the cleavage experiments in panels C and D; HJ substrate labeled in segments 1–4 were used in panels G and H. C. Hydroxyl radical cleavage of HisFlagMlh1-Mlh3 with FeBABE-labeled ssDNA (left) or HJ (right). Mlh1 cleavage fragments were detected by anti-Flag western blotting. Traces of individual lanes are shown. Experiments were done at least 3 times with comparable results. D. Hydroxyl radical cleavage of Mlh1-HisFlagMlh3 with FeBABE-labeled ssDNA (left) or HJ (right). Mlh3 cleavage fragments were detected by anti-Flag western blotting. E. Summary of the hydroxyl radical cleavage results. For each cleavage fragment of Mlh1 (top) and Mlh3 (bottom), a short stretch of amino acids in the vicinity of the predicted DNA-binding residues is indicated by a gray tick mark that is shaded in proportion to the intensity of the hydroxyl radical cleavage signal detected with each DNA substrate. The protein domains are cartooned along the horizontal axes: orange (Mlh1) or yellow (Mlh3) for linker regions; green for N and C-terminal domains of Mlh1; and blue for N and C-terminal domains of Mlh3. F. The ratio of cleavage fragment intensities mapping to the N-terminal domains and linkers of Mlh1 and Mlh3 with the ssDNA and HJ substrates. Error bars are standard deviations of three independent experiments. G, H. Effect of changing the position of the FeBABE moiety along the HJ substrate. (G) HisFlagMlh1-Mlh3 and (H) Mlh1-HisFlagMlh3. I. The positions of FeBABE-predicted DNA-binding residues are highlighted in red on a homology-based model of the Mlh1-Mlh3 heterodimer. J. Electrostatic potential map of the Mlh1-Mlh3 model.

The FeBABE positions along the ssDNA and HJ substrates are illustrated in **Fig 3A, B**. We assembled nucleoprotein complexes on immobilized FeBABE-conjugated DNA substrates using MutLγ complexes that carried an N-terminal tag either on Mlh1 (^HisFlag^Mlh1-Mlh3, **Fig 3C**) or Mlh3 (Mlh1-^HisFlag^Mlh3, **Fig 3D**). After the hydroxyl radical cleavage reaction, the samples were separated by SDS-PAGE and the cleavage fragments were detected by anti-Flag western blotting. The cleavage fragments were dependent on the presence of the FeBABE modification and activation with H_2_O_2_ (e.g., compare lane 2 with lanes 1, 3 and 4 in **Fig 3C**). The cleavage pattern was also specific to which protein was tagged and to the DNA substrate used (compare **Fig 3C** and **D**, left and right panels).

We mapped positions of the cleavages using molecular weight markers (see Materials and Methods). With the ssDNA substrate, the predicted cleavage positions map within the linker and the N-terminal domains of both Mlh1 and Mlh3 (**Fig 3C, D, E**). With the HJ substrate, the pattern was similar but the fragments corresponding to cleavage in the linkers were enriched relative to cleavages mapping to the N-terminus, as compared to the ssDNA substrate (**Fig 3F**). These findings suggest that the specificity for the HJ substrate relies more on the linker regions of Mlh1 and Mlh3 than does the specificity for the ssDNA substrate (**Fig 3E**).

We further asked whether the pattern of the hydroxyl radical cleavage fragments was affected by the position of the FeBABE moieties along the HJ substrate (**Fig 3G, H**). The intensity of the cleavage fragments decreased as the FeBABE probes were moved more than ~25 bp away from the center of the HJ substrate, indicating that the main protein-DNA contacts are located toward the center of the HJ (**Fig 3B, G, H**). However, the sizes of the cleavage fragments changed little if at all as the FeBABE positions of the probes were shifted. This may suggest that the structure and disposition of the Mlh1 and Mlh3 linker regions around the HJ core is not well defined and that the cleavage pattern results from a heterogenous mixture of protein/DNA complexes. Alternatively, it may reflect the limit of resolution of the footprinting strategy, e.g., because of the effective radius of hydroxyl radical damage and/or locations of protein segments that are particularly susceptible to cleavage.

We next sought to identify amino acid residues in the vicinity of the hydroxyl radical cleavage sites that might be directly involved in protein-DNA contacts. An alignment of fungal Mlh1 and Mlh3 proteins revealed conserved basic residues near each predicted cleavage site (**S1 Fig** and **Table 1**). When mapped to a homology-based model of the N- and C-terminal domains of the Mlh1-Mlh3 heterodimer, these candidate residues clustered to positively charged surfaces within the N-terminal domain, suggesting that they may indeed be part of a DNA-binding surface (compare position of the amino acids highlighted in red in **Fig 3I** to positive (blue) surfaces in **Fig 3J**). Predicted DNA-binding residues located within the linker regions of Mlh1 and Mlh3 could not be modeled because no structural information is available, but sequence alignments revealed that mapped cleavage sites are located within the most conserved regions of the linkers (**S1 Fig**).

**Table 1:**
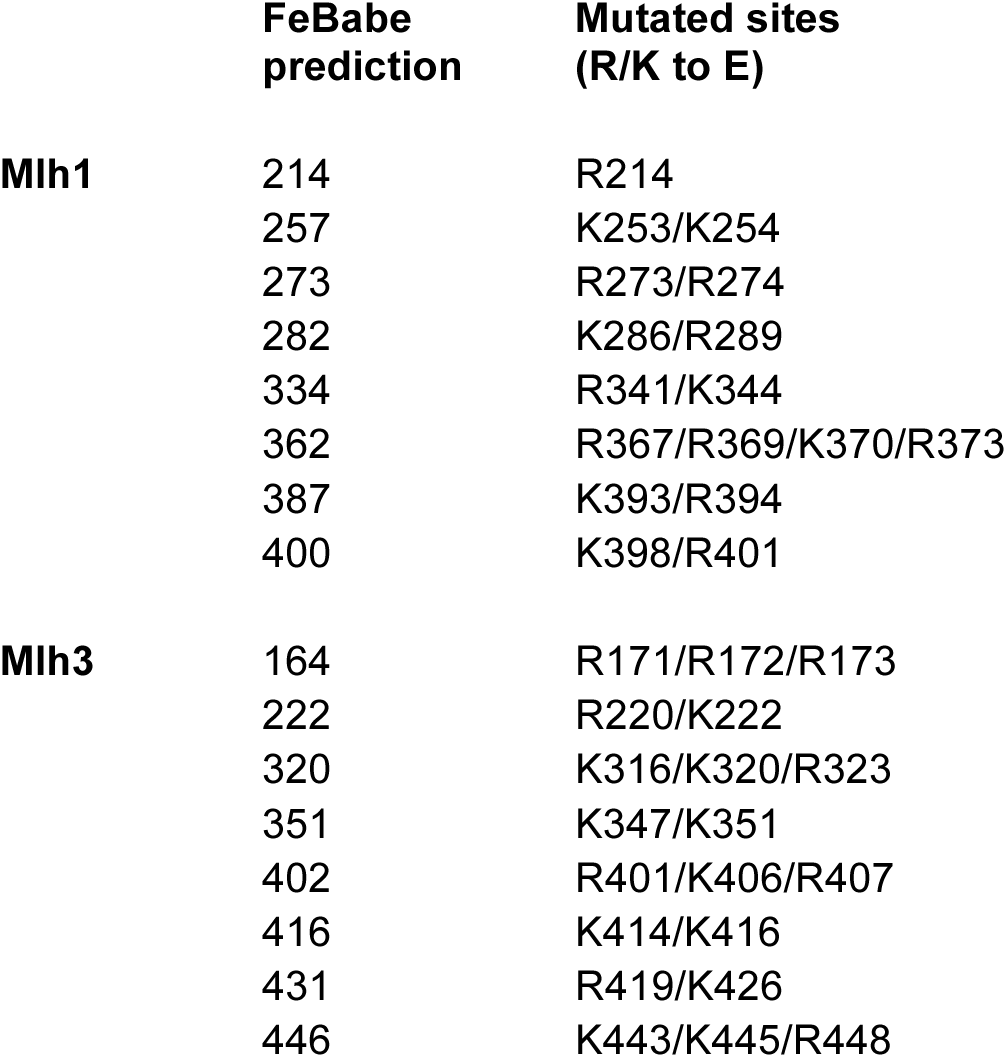
**Summary of the FeBABE-predicted DNA-binding sites.** The table shows the calculated hydroxyl radical cleavage positions and the corresponding amino acid mutations chosen for each predicted binding site.

For each subunit, we mutated eight sets of amino acids that corresponded to eight FeBABE cleavage sites mapped either in the N-terminal domain (N-terminal mutants) or in the flexible linker region (linker mutants) and tested the effects of these mutations on meiotic recombination and MMR *in vivo* (**Table 1**).

## Effects on meiotic recombination of mutating putative MutLγ DNA-binding residues

To address the roles of the predicted DNA-binding residues of Mlh1 and Mlh3 in meiotic recombination we introduced targeted mutations by gene replacement in yeast cells that harbor spore-autonomous fluorescent reporters. The mutant proteins were expressed as untagged versions from their endogenous promoters. The diploid cells had transgenes encoding red and green fluorescent proteins located next to the centromere and *ARG4* loci, respectively, of one chromosome VIII homolog, and blue fluorescent protein next to the *THR1* locus of the other homolog (**Fig 4A**). The transgenes are under the control of the *YKL050c* promoter, which is expressed late in sporulation, after meiosis, and therefore allows fluorescent proteins to be detected only in spores that inherit a copy of the reporter gene [57]. Upon sporulation, the frequency of crossing over in the two test intervals (*CEN8-ARG4* and *ARG4-THR1*) and the frequency of missegregation of chromosome VIII (MI nondisjunction) can be scored by the diagnostic fluorescence patterns of tetrads (**Fig 4A**).

**Fig 4:**
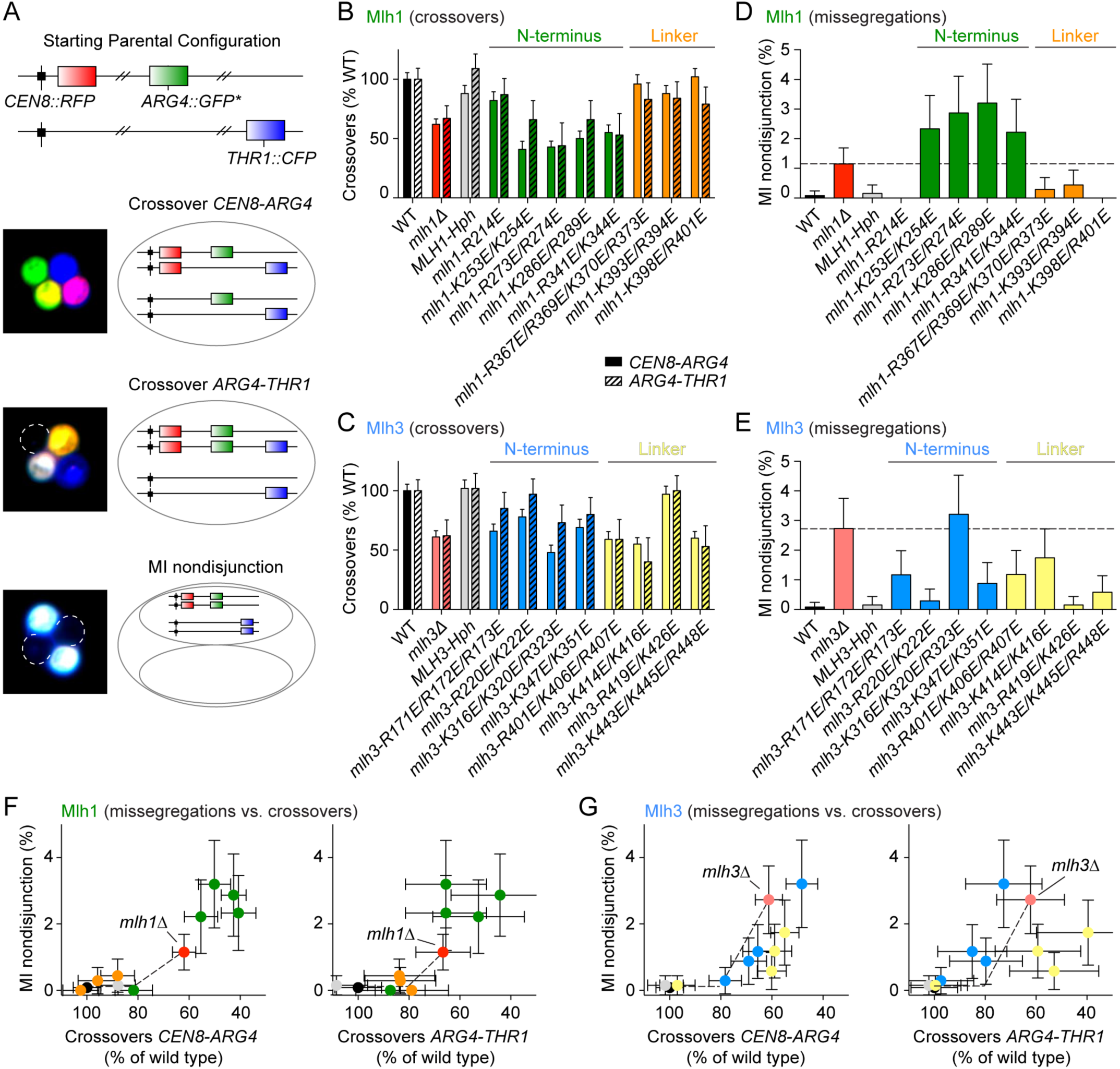
Effects of putative DNA-binding mutations of Mlh1 and Mlh3 on meiotic recombination and chromosome segregation. A. Spore-autonomous fluorescent reporter assay. The strains harbor fluorescent reporters at the *CEN8* and *ARG4* loci of one homolog of chromosome VIII and at the *THR1* locus of the other homolog. The fluorescence pattern of the tetrad allows detection of crossovers in the two test intervals (*CEN8 – ARG4* and *ARG4 – THR1*) and MI nondisjunction of chromosome VIII. Figure was adapted from [57]. B, C. Effects of the *mlh1* (B) and *mlh3* (C) mutations on crossover frequencies within two test intervals. The data are normalized to the crossover levels of the wild-type (WT) strain (CO*CEN8–ARG4* = 11.63 ± 0.63 cM; CO*ARG4–THR1* = 4.22 ± 0.39 cM). Error bars represent standard errors. D, E. Effects of the *mlh1* (D) and *mlh3* (E) mutations on MI nondisjunction of chromosome VIII. Error bars are 95% confidence intervals of the proportion. The dotted lines indicate the MI nondisjunction frequencies of the deletion strains. *MLH1-Hph*, *MLH3-Hph* and all putative DNAbinding mutant strains have a hygromycin resistance cassette integrated ~50 bp downstream of the corresponding gene. The data for *MLH1-Hph* and *MLH3-Hph* (gray) are controls to show that the hygromycin resistance cassette does not compromise the meiotic functions. F, G. Correlation between crossover frequency and MI nondisjunction in *mlh1* (F) and *mlh3* (G) mutants. Crossover frequencies in the test intervals *CEN8-ARG4* and *ARG4-THR1* are expressed as a percent of crossovers in wild type. The x axes are plotted so that the crossover defect increases from left to right. On the y axes, the defects in chromosome segregation (expressed as percent of cells showing MI nondisjunction of chromosome VIII) increase from bottom to top. The color code is as in **panels B-E.**

Crossover patterns can be scored unambiguously. However, the MI nondisjunction pattern is ambiguous because it can also arise from a four-chromatid double crossover in the *ARG4-THR1* interval. In the wild type, this segregation pattern is seen at a frequency of about ~0.1% (1 out of 1268 tetrads). Double crossovers in the *CEN8-ARG4* interval, which can be scored unambiguously, were very rare in all strains (<0.05 to 0.2%). Because the genetic interval of *ARG4-THR1* (4.2 ± 0.39 cM) is smaller than *CEN8-ARG4* (11.63 ± 0.63 cM), double crossovers in this interval should be even less frequent. Using a separate assay that does not suffer from this ambiguity, Thacker *et al.* found that the MI nondisjunction frequency of chromosome VIII in wild type is ~0.1% [57]. Since this is the frequency at which we detected the ambiguous pattern in wild type, we treated all of these events as MI nondisjunction. We may have therefore slightly underestimated the crossover frequency in the *ARG4-THR1* interval in wild type, but this is unlikely to be a significant bias in crossover-deficient strains for which MI nondisjunction dominates.

We scored over 600 tetrads per strain (**Fig 4** and see **S1 Table** for genetic distances and statistical analyses). The crossover frequency in *mlh1Δ* and *mlh3Δ* strains was reduced to about 60–70% of the wild-type values in the two intervals, and the residual crossover levels were not significantly different between these strains (p > 0.9, G test), in agreement with prior studies [14, 35, 49] (**Fig 4B, C**). MI nondisjunction frequency increased from ~0.1% in wild type to 1.15% ± 0.54% in *mlh1Δ* and 2.73% ± 1.02% in *mlh3Δ* (mean ± 95% CI)(**Fig 4D, E**). This 2.4-fold difference was statistically significant (p = 0.005, Fisher’s exact test) and to our knowledge has not been reported previously.

Most of the *mlh1* mutations affecting the N-terminal domain resulted in a significant decrease in crossing over and an increase in MI nondisjunction (green bars, **Fig 4B, D**). Interestingly, at least three of the *mlh1* N-terminal domain mutants (*mlh1-K253E/K254E*, *mlh1 R273E/R274E* and *mlh1-K286E/R289E*) showed a stronger phenotype than the *mlh1Δ* strain, particularly for MI nondisjunction (green bars, **Fig 4B, D**). In contrast, the *mlh1* linker mutants (*mlh1-R367E/R369E/K370E/R373E*, *mlh1-K393E/R394E* and *mlh1-K398E/R401E*) conferred little or no meiotic defect (orange bars, **Fig 4B, D**).

The *mlh3* mutants were more variable: some had weak defects and there was no clear distinction between linker mutants and N-terminal mutants, unlike for *mlh1*. For example, in contrast to *mlh1*, three of the *mlh3* linker mutants showed a significant meiotic defect (yellow bars, **Fig 4C, E**) and none of the *mlh3* mutants had a significantly stronger phenotype than the *mlh3Δ* strain.

When MI nondisjunction frequency was plotted as a function of crossover level in either test interval, MI nondisjunction was increased only in those mutants where the crossover frequency had dropped below ~70–80% of wild-type levels (**Fig 4F, G**). This agrees with a previous observation that spore viability only starts to decrease below a certain threshold of crossovers [58]. We also found that *mlh3* linker mutants exhibited a proportionally stronger defect in crossing over than in chromosome segregation compared to the *mlh3Δ* strain (compare yellow points to the red point in **Fig 4G**).

### Effects on MMR of mutating putative MutLγ DNA-binding residues

We used a Lys+ reversion assay to quantify the effects of the predicted DNA-binding mutations in MMR (**Fig 5** and **S2 Table**). The assay takes advantage of a mutant *lys2* gene containing a stretch of 14 A residues that creates a hotspot for DNA polymerase slippage [59]. The allele has a +1 bp frameshift, so changes of −1 bp or +2 bp can give Lys+ revertants. MMR-deficient strains such as *mlh1Δ* frequently fail to repair mutations introduced by DNA polymerase, causing elevated Lys+ reversion frequencies.

**Fig 5:**
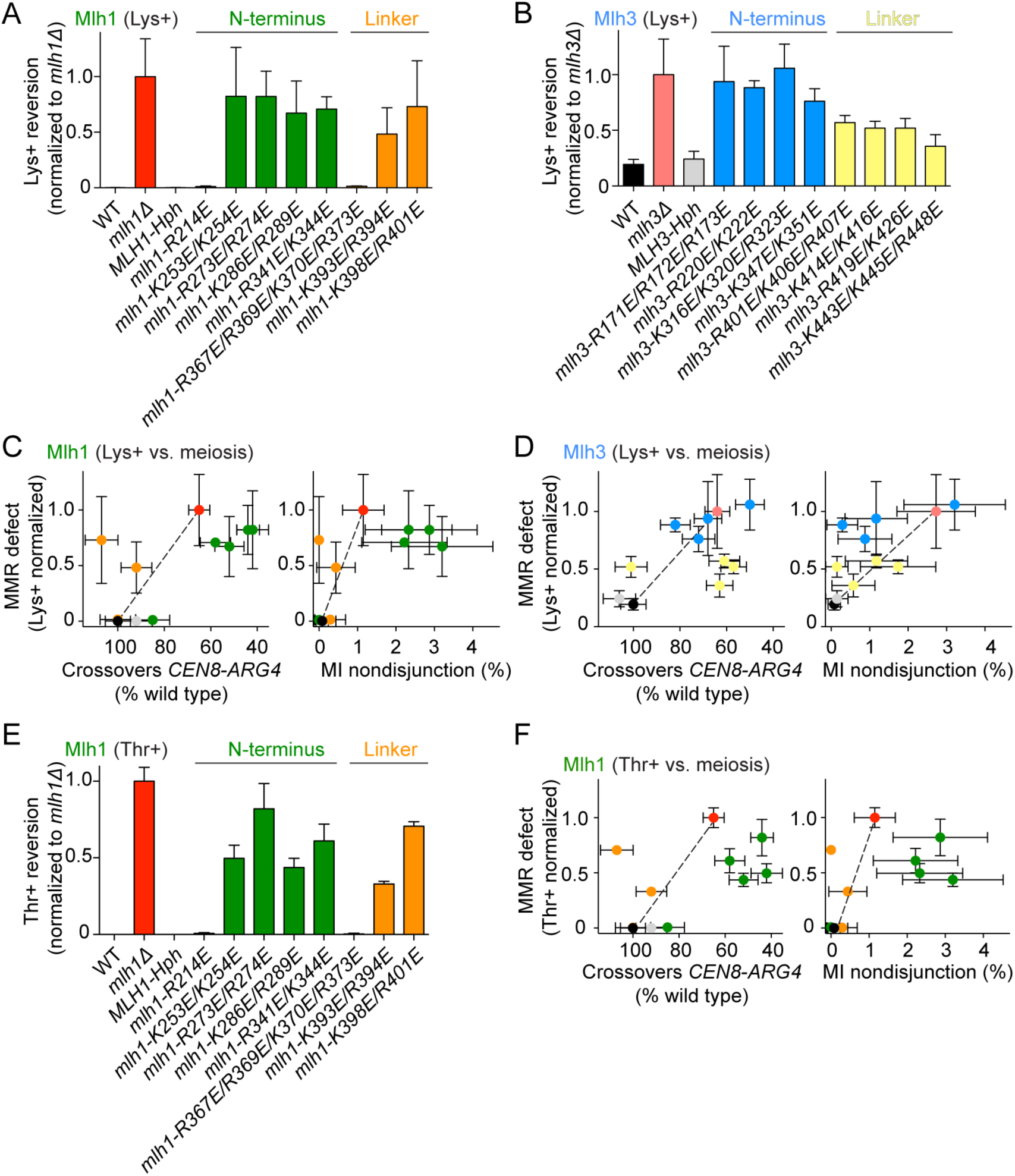
Effects of putative DNA-binding mutations of Mlh1 and Mlh3 on mismatch repair. A. Lys+ reversion in *mlh1* mutants, normalized to the reversion frequency of the *mlh1*strain. Error bars are standard deviations from 3–6 independent cultures. Lys+ reversion frequency of wild type was < 10-4 and *mlh1*was 4 10-2. B. Lys+ reversion in *mlh3* mutants, normalized to the reversion frequency of the *mlh3*strain. Error bars are standard deviations from 3–6 independent cultures. Lys+ reversion frequency of *mlh3*was 1.7 10-4. C, D. Correlations between MMR defect and either the crossover defect (left) or MI nondisjunction (right) in *mlh1* (C) and *mlh3* (D) mutants. The x axes are plotted so that the crossover defect increases from left to right. The y axes are Lys+ reversion frequencies normalized to the deletion strains as in panels A and B. The defect increases from the bottom to top of the graph. The color code is as in **panels A, B** and **Fig 4.** Mutations within the same domain tend to cluster together, reflecting the relative importance of the N-terminal domain and linker regions in meiotic recombination and MMR. Points that deviate from the diagonals (dotted lines) that link the wild-type strain to *mlh1*or *mlh3*are separation-of-function mutants. E. The frequency of Thr+ reversion in *mlh1* mutants, normalized to the reversion frequency in the *mlh1*strain. Error bars are standard deviations from three independent cultures. Thr+ reversion frequency of wild type was < 10-6 and *mlh1*was 10-4. F. Correlations between MMR defect in the Thr+ assay and either the crossover defect (left) or MI nondisjunction (right) for *mlh1* mutants.

With this sensitive assay, the Lys+ reversion frequency of an *mlh1Δ* strain was three orders of magnitude higher than wild-type levels (**Fig 5A** and [60]). Out of the eight *mlh1* point mutants we examined, two were nearly indistinguishable from a wild-type stain (*mlh1-R214E* and *mlh1-R367E/R369E/K370E/R373E*), whereas the other six mutants reached Lys+ reversion levels close to *mlh1Δ*.

Because Mlh3 plays only a minor role in MMR, the Lys+ reversion frequency of an *mlh3Δ* strain was only ~5-fold greater than in wild type (**Fig 5B** and [33, 49]). All eight *mlh3* mutants conferred a significant MMR defect, with the N-terminal mutants almost indistinguishable from *mlh3Δ* (**Fig 5B** and **S2 Table**).

When the MMR defects were plotted versus crossover defects or chromosome missegregation frequencies, two patterns emerged. First, the mutants clustered by class on the basis of mutation location in either the N-terminal domain or in the linker region, indicating that different contributions in MMR or meiotic recombination can be attributed to specific actions of particular domains. Second, several mutants substantially deviated from the diagonals formed between wild type and the deletion strains; these are therefore separation-of-function mutants (**Fig 5C, D**). Specifically, *mlh1* linker mutants (particularly *mlh1-K398E/R401E*) were above the diagonal, consistent with the possibility that DNA binding by the Mlh1 linker may be more important for MMR than meiotic recombination (orange dots, **Fig 5C** left and right panels). The *mlh1* N-terminal mutants clustered right of the diagonal, reflecting their stronger defect in meiotic recombination than *mlh1Δ* (green dots, **Fig 5C** left and right panels).

The *mlh3* mutants also tended to cluster according to mutation position (**Fig 5D**). When the MMR defect was plotted against crossovers, three of the *mlh3* linker mutants (yellow points; *mlh3-R401E/K406E/R407E*, *mlh3-K414E/K416E* and *mlh3-K443E/K445E/R448E*) clustered below the diagonal, whereas the N-terminal mutants (blue points) fell close to the diagonal. When MMR defect was plotted against MI nondisjunction, however, the linker mutants fell on the diagonal while N-terminal mutants lay above the diagonal. Thus, the *mlh3* linker mutants confer a disproportionately stronger defect in crossing over than in either meiotic chromosome segregation or MMR.

The 14-A insertion Lys+ reversion assay was chosen because it allowed us to score mutations in either *mlh1* or *mlh3*, but it also yields high basal rates in wild type [49, 59]. Therefore, we also measured the MMR deficiency of the *mlh1* mutants in an independent Thr+ reversion assay that uses a +1 T insertion in a stretch of 6 T’s in the *HOM3* gene (*hom3-10*) [61]. *mlh3* mutants were not scored because the Thr+ reversion rate of *mlh3Δ* was not detectably higher than background. Levels of Thr+ reversion in the *mlh1* mutants, when normalized to *mlh1Δ*, were similar to Lys+ reversion (**Fig 5E,** compare with **5A**). A comparison of MMR defects in the Thr+ reversion assay with meiotic recombination defects revealed similar clustering of the *mlh1* mutant classes as with the Lys+ reversion assay, thus our conclusions are robust to differences between these experimental systems (**Fig 5F**).

### *In vitro* characterization of DNA-binding mutants

We went on to characterize a selection of MutLγ mutants *in vitro*. For both Mlh1 and Mlh3, we chose one N-terminal mutant (*mlh1-K286E/R289E* and *mlh3-K316E/K320E/R323E*) and one linker mutant (*mlh1-K393E/R394E* and *mlh3-K414E/K416E*) and purified single and double mutant complexes (**Fig 6A**). By EMSA with ssDNA or HJ substrates, all of the mutants displayed binding defects although all retained some DNA-binding activity (**Fig 6B-E**). In addition, the double mutants were significantly more affected than either single mutant.

**Fig 6:**
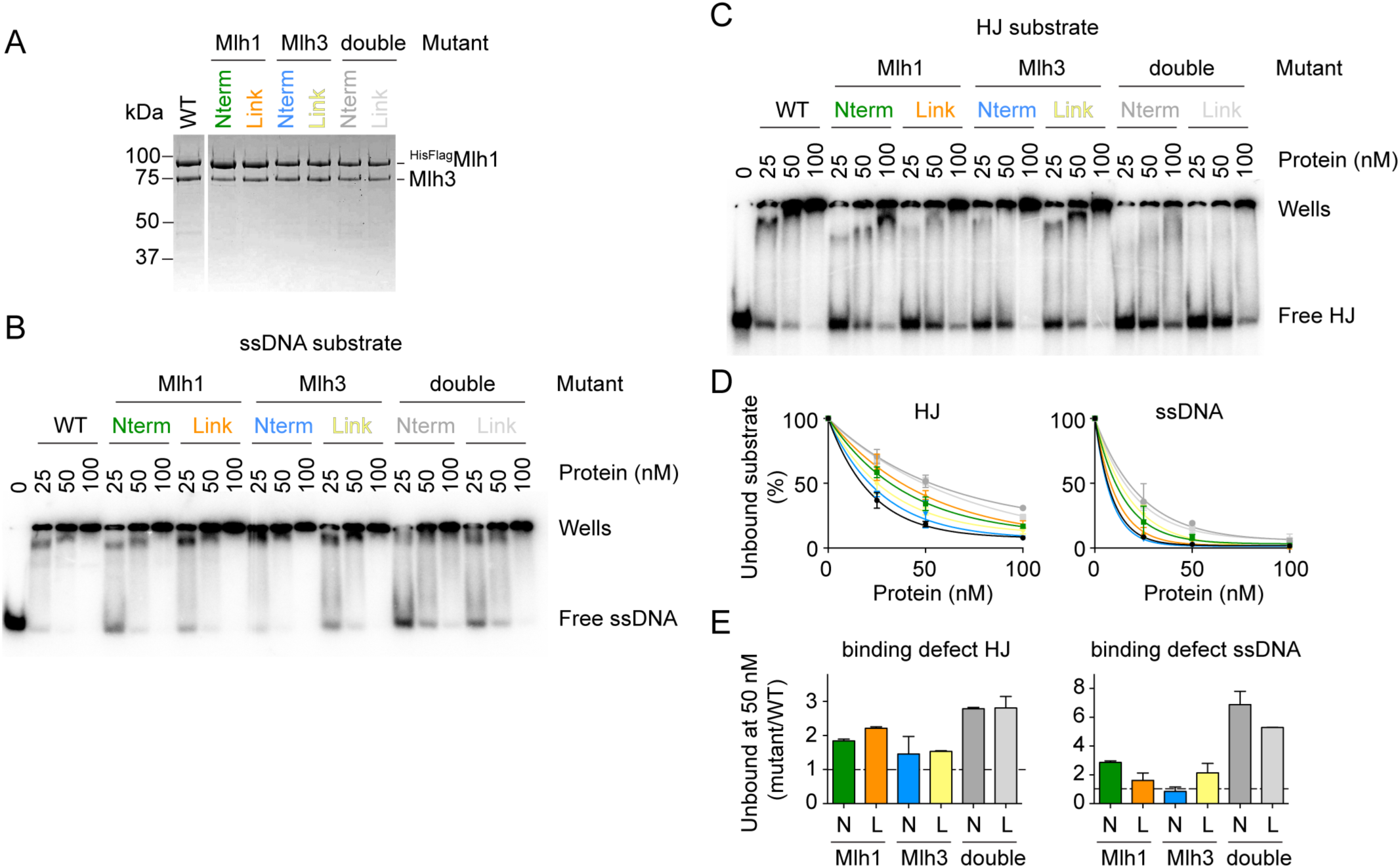
*In vitro* characterization of a selection of MutLγDNA-binding mutants. A. Purification of MutLγDNA-binding mutants. All complexes carried an N-terminal tag on Mlh1. The mutants are as follows: Mlh1-Nterm (K286E/R289E), Mlh1-Link (K393E/R394E) and Mlh3 - Nterm (K316E/K320E/R323E), Mlh3-Link (K414E/K416E). A Coomassie-stained SDS-PAGE gel is shown. B. EMSA with MutLγmutants using 1 nM end-labeled ssDNA substrate. C. EMSA of MutLγmutants with 1 nM end-labeled HJ substrate. D. Quantifications of the free substrate in EMSA experiments from panels B and C. Error bars show the range from two independent experiments. E. The HJ and ssDNA-binding defects of the MutLγmutants measured by the ratio of free substrate with mutant MutLγdivided by the free substrate with wild-type MutLγat 50 nM protein. Error bars show the range from two independent experiments.

The degree to which the ssDNA or HJ binding activities of each mutant were compromised varied. We asked whether these could be predicted from the intensity of the corresponding FeBABE-induced cleavage signal, but this was not straightforward. For example, the Mlh1 linker mutant had a stronger defect in HJ binding than the N-terminal mutant (**Fig 6E**). This is consistent with the hydroxyl radical footprinting result because the cleavage fragments with the HJ substrate mapped to the linker and not to the N-terminal domain (see **Fig 3F**). In the case of Mlh3, the difference in HJ binding defect between the N-terminal and linker mutants was not obvious (**Fig 6E**). This is explained by the fact that, in contrast to the *mlh1* N-terminal mutant, the HJ substrate had significant hydroxyl radical cleavage signal that mapped to the region of the *mlh3* N-terminal domain that we mutated (*mlh3-K316E/K320E/R323E*) (see **Fig 3F**). Hence, a difference in HJ binding affinity between the purified Mlh3 linker and N-terminal mutant complexes was not necessarily expected.

In the case of the ssDNA binding activity, the mapped DNA-binding interface is extensive (**Fig 3**) and no clear predictions can be made as to the effect of a specific DNA-binding mutation. Indeed, the FeBABE footprinting assay establishes proximity, not direct binding, so there is no *a priori* way of predicting which residues are the most critical for a specific binding activity based on hydroxyl radical cleavage intensity alone. Taken together, the DNA-binding defects of the MutLγ mutants show that the linker and N-terminal domains exert different contributions to the specificity of the DNA substrate. The binding defects globally agreed with the predictions of the hydroxyl radical footprinting assay, as far as it was possible to make clear predictions.

## Meiotic phenotypes combining mutations of MutLγ with other recombination mutations

To better understand the function of MutLγ in meiotic recombination and the effects of the DNA-binding mutants, we tested the effect of the *mlh1* and *mlh3* mutations in the context of *mms4Δ* or *msh5Δ* backgrounds (**Fig 7** and **s2 Table**). Mms4 is a component, together with Mus81, of a structure-selective nuclease; it contributes to a quantitatively minor crossover pathway in wild type but is believed to resolve the excess of joint molecules that accumulate when the MutLγ pathway is compromised [1, 17, 22, 28]. Msh5 is part of the MutSγ complex, which channels recombination intermediates toward the MutLγ-dependent crossover pathway [1, 12, 18].

**Fig 7:**
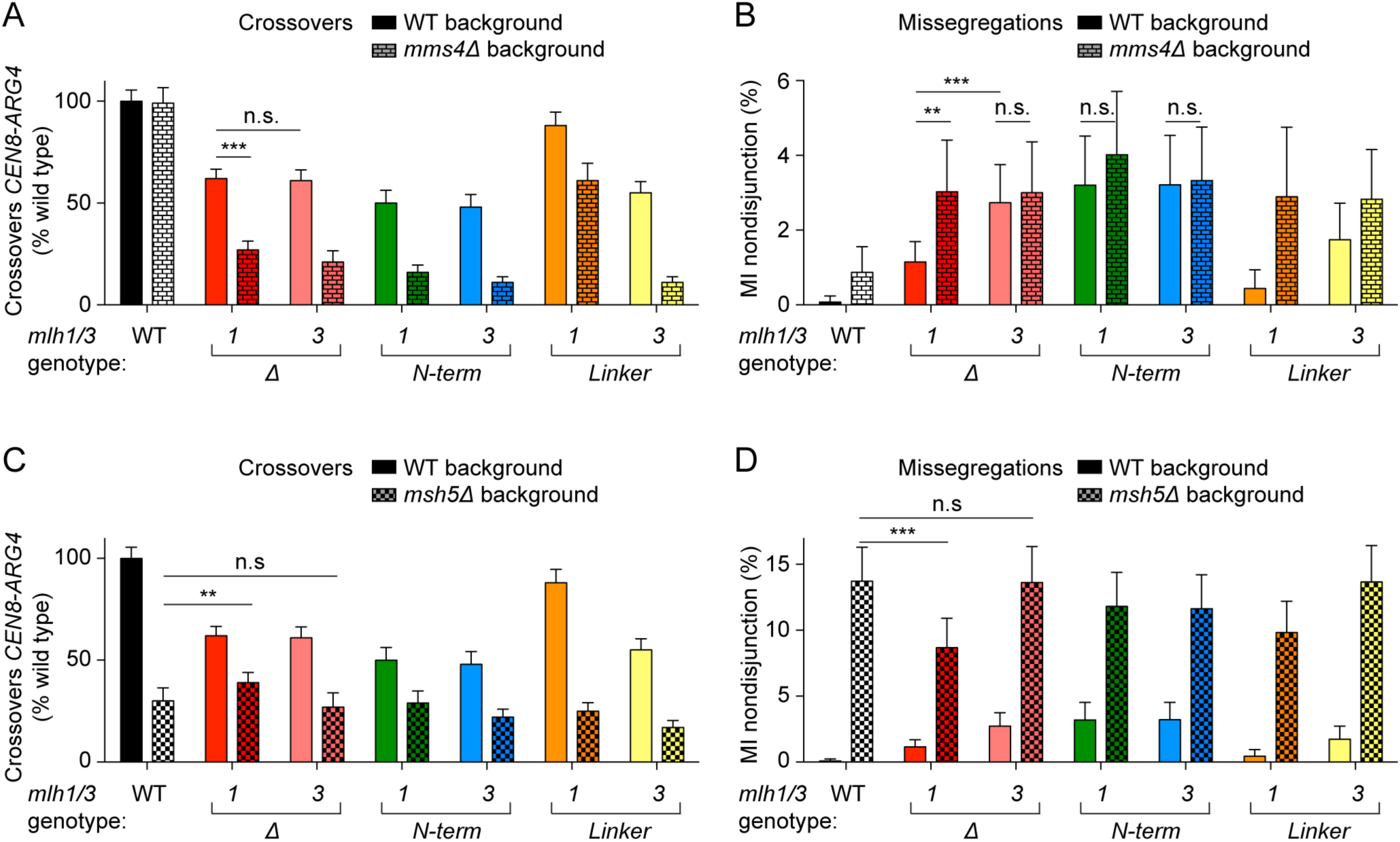
Effects on meiotic recombination of MutLγmutations in *mms4*and *msh5*backgrounds. Crossover levels within the *CEN8-ARG4* interval and chromosome VIII MI nondisjunction frequencies were measured by the fluorescent spore assay. The data of MutLγmutants in a wild-type background are reproduced from **Fig 4** to facilitate comparison. The *mlh1* and *mlh3*mutants assayed were: *mlh1-Nterm* (*K286E/R289E*), *mlh3-Nterm* (*K316E/K320E/R323E*), *mlh1-Link* (*K393E/R394E*) *mlh3-Link* (*K414E/K416E*). A. Crossover levels of MutLγmutants in wild-type and *mms4*backgrounds. Error bars represent standard errors. The data were normalized to crossover levels in wild-type (11.63 ± 0.63 cM). B. MI nondisjunction frequencies of MutLγmutants in wild-type and *mms4*backgrounds. Error bars are 95% confidence intervals of the proportion. C. Crossover levels of MutLγmutants in wild-type and *msh5*backgrounds. D. MI nondisjunction frequencies of MutLγmutants in wild-type and *msh5*backgrounds. Statistically significant differences are indicated with asterisks (**, p<0.01; ***, p<0.001). Selected non-significant differences are highlighted (n.s.).

<u>**Epistasis with *mms4Δ:***</u> Deletion of *MMS4* had no detectable effect on crossover frequency in the *CEN8-ARG4* interval (**Fig 7A**). However, MI nondisjunction was increased from ~0.1% in wild type to ~1% in *mms4Δ* (**Fig 7B**). Crossovers in *mms4Δ mlh1Δ* were ~30% of wild-type levels and MI nondisjunction increased to ~3% (**Fig 7A, B**), in agreement with previous reports showing that *mms4Δ* and *mlh1Δ* are not epistatic [17, 23]. *mlh3Δ* behaved like *mlh1Δ* in terms of crossing over, thus *mlh3Δ* is also not epistatic with *mms4Δ* (**Fig 7A**). However, the double mutant *mms4Δ mlh3Δ* had similar levels of MI nondisjunction as *mlh3Δ* and *mms4Δ mlh1Δ* (**Fig 7B**). Thus, *mlh3Δ* is epistatic to *mms4Δ* in terms of MI nondisjunction.

The *mlh1* and *mlh3* N-terminal mutants (*mlh1-K286E/R289E* and *mlh3-K316E/K320E/R323E*) behaved similarly to *mlh3Δ* in that their high levels of MI nondisjunction were not further increased when combined with *mms4Δ* (**Fig 7B**). The *mlh1* linker mutant (*mlh1-K393E/R394E*), which had a mild phenotype in a wild-type background, showed a significantly worse defect in a *mms4Δ* background, both in terms of crossovers and MI nondisjunction (**Fig 7A, B**). The *mlh3* linker mutant (*mlh3-K414E/K416E*) had an intermediate phenotype: It behaved like *mlh3Δ* in terms of crossing over, but MI nondisjunction in the single mutant was lower than with *mlh3Δ* and reached the levels of *mlh3Δ* when combined with *mms4Δ* (**Fig 7A, B**).

In summary, none of the mutants were epistatic with *mms4Δ* in terms of crossing over, as expected. However, *mlh3Δ* and the *mlh1* and *mlh3* N-terminal mutants (but not *mlh1Δ*) were epistatic to *mms4Δ* for MI nondisjunction. Implications of these findings are addressed in the Discussion.

<u>**Epistasis with *msh5Δ:***</u> Current models of meiotic recombination places MutSγ upstream of MutLγ, with MutSγ proposed to stabilize recombination intermediates and recruit MutLγ for crossover formation [1, 18]. A straightforward expectation, therefore, would be that absence of MutSγ would be epistatic to mutations in MutLγ. To test this, we asked whether the different phenotypes that we observed between *mlh1* and *mlh3* mutants would reveal different behaviors in the context of *msh5Δ* mutants.

Deletion of *MSH5* decreased crossovers in the *CEN8-ARG4* interval to ~30% of wild-type levels and increased MI nondisjunction to ~13.7%, consistent with published reports (**Fig 7C, D**) [58, 62]. Surprisingly, when the *mlh1Δ* mutation was combined with *msh5Δ*, the MI nondisjunction frequency was decreased compared to the *msh5Δ* single mutant (to ~8.7%, p =0.0047, Fisher’s exact test) (**Fig 7D**). In contrast, the MI nondisjunction frequency of *mlh3Δ msh5Δ* was similar to *msh5Δ*. Thus, *mlh1Δ* but not *mlh3Δ* can partially alleviate the chromosome missegregation defect of *msh5Δ*. In addition, *mlh1Δ* also partially rescued the crossover defect of *msh5Δ*. The effect was quantitatively small (from 30.1% of wild-type crossover levels in *msh5Δ* to 38.7% in *mlh1Δ msh5Δ*) but statistically significant (p = 0.01, G test). Here, too, *mlh3Δ* had no effect on the phenotype of *msh5Δ*.

The *msh5Δ* mutation was epistatic to most of the tested *mlh1* and *mlh3* DNA-binding mutations for both crossing over (**Fig 7C**) and MI nondisjunction (**Fig 7D**), except that the *mlh1* linker mutant *(mlh1-K393E/R394E*) also partially rescued the MI nondisjunction frequency of *msh5Δ* (p = 0.038, Fisher’s exact test).

Thus, *mlh1Δ* and to a lesser extent the *mlh1* linker mutant partially rescued the crossover and MI nondisjunction phenotype of *msh5Δ*. This behavior is not explained by current models in which MutSγ and MutLγ follow a strictly linear genetic relationship. Possible reasons for this behavior are discussed below.

## Discussion

MutL-family complexes exhibit different activities, including DNA binding and cleavage, ATP binding and hydrolysis, conformational transitions, and functional interactions with other protein factors [31, 50]. For MutLα, interdependence of these activities has been partially elucidated *invitro*: for example, DNA binding by MutLα depends on ATP binding and nuclease activity is regulated by protein interactions. Nevertheless, the detailed mechanism of MutLα in MMR is still unclear [31, 50]. For MutLγ, genetic experiments have established that ATP binding and/or hydrolysis, DNA binding, DNA cleavage, and protein-protein interactions (e.g., with Exo1) are all required for meiotic crossover formation (this work and [17, 19, 47, 49, 58, 61, 63]). However, *invitro* experiments have so far been unable to demonstrate the dependence of any of these activities on one another, except for ATP-driven conformational changes (this work). For example, no effects of ATP on DNA-binding and cleavage or effects of DNA on the ATPase activity were detected, suggesting that the physiologically relevant activities have not been fully reconstituted (this work and [45, 46]).

Here, biochemical characterization revealed unexpected properties of MutLγ regarding its specificity for DNA substrates, the contributions of different DNA-binding surfaces, and the assembly of nucleoprotein complexes into higher-order structures. We also generated a collection of MutLγ DNA-binding mutants that are defective in MMR and/or meiotic recombination, supporting the conclusion that DNA binding is critical for MutLγ function. Indeed, although DNA binding, including to ssDNA, was reported for MutLα and MutL, a role for DNA binding in MMR has remained controversial for decades [51, 55, 64–66]. The strongest evidence for MutLα comes from a DNA-binding defective mutant that retains all of its DNA-independent activities *in vitro* but behaves like a null mutant *in vivo* [65]. Our findings show that this is also the case for MutLγ and further indicate that distinct DNA-binding surfaces of the complex are associated with distinct specificities for DNA substrates and exert separable biological functions.

### Specificity for ssDNA and HJ substrates and overlapping DNA-binding sites

MutLγ recognition of HJs seems easy to rationalize given that the enzyme is proposed to resolve dHJs into crossovers [1, 18]. However, binding to ssDNA is more surprising because canonical models of meiotic recombination do not predict that ssDNA will be present at the stage of the dHJ intermediate. Although the DNA-binding activity of MutLγ is not particularly strong (in the two-digit nM range), the affinity for ssDNA is considerably higher than for HJs under our experimental conditions. One possibility is that MutLγ plays a role earlier in recombination than typically envisioned, for example, by binding to ssDNA exposed at nascent strand-exchange events.

Another surprising observation was that the HJ and ssDNA binding activities are separable — i.e., largely non-competing — implying that the binding sites must be different and raising the possibility that the complex can bind two substrates simultaneously. FeBABE footprinting helped explain this observation by revealing that the DNA-binding interfaces for the two substrates only partially overlap. The interface for HJ binding appears to lie primarily within the linker regions of Mlh1 and Mlh3, while the ssDNA-binding interface appears more extensive and includes both the N-terminal domains and the linker regions.

What might be the role of these overlapping binding sites? Perhaps affinity for ssDNA and simultaneous binding to different substrates assists in recruiting MutLγ to recombination sites. During meiotic recombination, there are thought to be two main stages where ssDNA is generated: during exonucleolytic resection of the double-strand breaks and during invasion of the homologous template, which exposes ssDNA from the template in the form of D-loops. We propose that MutLγ may be recruited to these early recombination intermediates, prior to formation of a dHJ. In this model, MutLγ may bind initially to ssDNA and then probe for nearby branched intermediates.

### DNA binding cooperativity and higher-order complexes

Yeast MutLα binds DNA cooperatively and prefers long substrates, and AFM experiments have shown that MutLα forms long tracks along DNA and that several substrates can be bound within the same higher order complex [55]. We found that MutLγ also binds DNA cooperatively, forms higher order structures, has a preference for longer DNA, and can bind multiple substrates simultaneously. It is currently unclear which, if any, of these higher order complexes are physiologically relevant. Interestingly, however, MLH1 and MLH3 form bright immunostaining foci that mark presumed crossover sites in mammalian cells [39, 40, 67, 68], suggesting that formation of higher order complexes is part of the crossover-promoting function of MutLγ. It is tempting to speculate that the oligomeric MutLγ-DNA complexes observed *in vitro* may be related to these foci.

### Contributions of putative DNA binding domains to MMR and meiotic recombination

To better understand how DNA binding relates to MutLγ function *in vivo*, we mutated residues that appear to be in proximity to DNA, as judged from hydroxyl radical footprinting. We consider it likely that at least some of these residues lie within protein surfaces that contact DNA directly. In support of this inference, the subset of mutants tested *in vitro* had diminished DNA binding.

Most *mlh1* N-terminal mutants showed decreased crossing over and increased missegregation of chromosome VIII, and were also defective in MMR. The MMR defects were comparable to *mlh1Δ*, but the meiotic defects were unexpectedly more severe than in the null (discussed further below). In contrast, the *mlh1* linker mutants showed little or no meiotic defect but were severely defective in MMR.

For Mlh3, the N-terminal mutants had variable degrees of meiotic defects but were more uniformly defective in MMR. In contrast, most of the *mlh3* linker mutants were defective in meiosis, but tended to show greater defects in crossing over than in chromosome segregation. Most *mlh3* linker mutants also conferred a partial MMR defect, but interestingly, the allele that was normal for crossing over (*mlh3-R419E/K426E*) was not the same as the allele that was nearly normal in MMR (*mlh3-K443E/K445E/R448E*).

Together, these findings indicate that these putative DNA-binding surfaces contribute to functions of both Mlh1 and Mlh3 in both MMR and meiotic recombination. The observed defects are consistent with DNA binding *per se* being important, but it is also possible that the mutations cause defects *in vivo* because of biochemical defects that are different from (or in addition to) DNA binding alterations. For example, the mutations might impinge on protein-protein interactions with crossover– or MMR-promoting factors such as Exo1 [63]. Importantly, identification of separation-of-function mutations in both proteins indicates that the molecular roles of Mlh1 and Mlh3 can be distinguished in MMR vs. meiotic recombination. Specifically, the Mlh1 linker region appears to be more important in MMR, while the Mlh3 linker region appears to be more important for meiotic crossover formation.

Several of the mutants were also analyzed in prior studies [47, 61, 69]. Mlh1– R273E/R274E had compromised DNA-binding activity for the Mlh1 N-terminal domain [69]. In addition, *mlh1* linker mutations conferred MMR defects (*K393A/R394A* and *R401A/D403A*) and N-terminal domain mutations conferred MMR and meiotic recombination defects (*K253A/K254A*, *R273A/R274A*) [61]. The Alani group recently characterized a collection of *mlh3Δ* mutants and also identified alleles with separable functions in MMR and crossing over [70]. Several of their mutants were similar to the ones reported here, including for example a *mlh3Δ* linker mutant (*mlh3-K414A/K416A*, referred to as *mlh3-32*), which conferred a stronger defect in crossing over than in MMR. Our observations agree well with these independent studies.

*In vitro* experiments with truncations of the Mlh1 and Pms1 linker domains previously implicated the linker regions of MutLα in DNA binding [71]. Here, we identified specific substitutions within the most conserved regions of the linkers of Mlh1 and Mlh3 that compromise the DNA binding activity (residues ~393-418 in Mlh1 and ~401-448 in Mlh3).

### Interplay between alternative crossover-forming pathways

Unlike MutLγ, the Mlh1-containing MutLα and MutL_β_ heterodimeric complexes are dispensable for meiotic crossing over, although they do participate in mismatch correction and other aspects of gene conversion during recombination [1, 18, 35]. Moreover, crossing over plays a critical role in promoting chromosome segregation, but gene conversion does not. Therefore, when we began this work, we envisioned that absence of either Mlh1 or Mlh3 would cause indistinguishable defects in crossover formation and therefore in chromosome segregation. However, we found that the meiotic phenotypes of the *mlh1Δ* and *mlh3Δ* strains differed: the mutants had similarly decreased crossover levels, but MI nondisjunction of chromosome VIII was 2- to 3-fold more frequent in *mlh3Δ*. This difference was not observed in previous measures of spore viability, which have been reported to be reduced to similar extents in the two mutants (to ~70–80%, e.g. [14, 19]). One possible reason for the apparent discrepancy with spore viability measurements could be chromosome-specific effects, since we assayed missegregation of only chromosome VIII, whereas spore viability reflects segregation of all chromosomes and is probably driven principally by behaviors of the smallest chromosomes. Another possibility is that the greater MMR defect in *mlh1Δ* results in a higher burden of haplolethal mutations accumulating during premeiotic growth, in turn yielding more inviable spores than accounted for by chromosome missegregation alone [14]. The spore-autonomous fluorescent reporter assay is independent of spore viability [57], so it should not suffer from this complication.

We further found that *mlh1* N-terminal mutants were worse than *mlh1Δ* for both crossing over and chromosome missegregation. We also identified instances where decreases in crossing over did not correlate closely with increases in MI nondisjunction. For example, *mlh3Δ* linker mutants (*mlh3-R401E/K406E/R407E*, *mlh3-K414E/K416E* and *mlh3 - K443E/K445E/R448E*) had crossover defects as strong as in *mlh3Δ* but showed less MI nondisjunction than *mlh3Δ*.

In summary, *mlh3* linker mutants behaved similarly to *mlh1Δ* (strongly reduced crossovers but only moderate MI nondisjunction), whereas *mlh1* N-terminal mutants were more similar to *mlh3Δ* (strongly reduced crossovers and high MI nondisjunction). The *mlh1* and *mlh3* N-terminal mutants were also similar to *mlh3Δ* in that they were epistatic to *mms4Δ* for MI nondisjunction but were non-epistatic with *mms4Δ* for crossing over.

Taken together, these results further illuminate the ways in which Mlh1 and Mlh3 contribute differently to meiotic recombination. For the null *mlh1Δ* and *mlh3Δ* mutations, differences may reflect fundamental distinctions between the proteins in terms of the complexes they inhabit: Mlh1 is part of MutLα, MutL_β_, and MutLγ, whereas Mlh3 is only known to be a component of MutLγ. Thus, the *mlh1Δ* phenotype may reflect absence of Mlh1 itself; absence of all three heterodimeric complexes (MutL_α_, MutL_β_, and MutLγ); aberrant behaviors of Mlh2, Mlh3, and Pms1 in the absence of their binding partner; or some combination of these defects. In contrast, the *mlh3Δ* phenotype could arise from absence of a specific function of Mlh3 itself; absence of MutLγ; and/or aberrant behavior of MutL_α_ or MutL_β_ caused by there no longer being competition between Mlh3 and other proteins for binding to Mlh1. Interestingly, MutL_β_ has recently been implicated in regulating meiotic gene conversion tract lengths [36], raising the possibility that differences between *mlh1Δ* and *mlh3Δ* could trace in part to differences in activity of MutL_β_.

Two main scenarios can be envisioned to explain why the mutants have different nondisjunction levels despite roughly similar crossover defects: MI chromosome segregation may be more accurate than expected given the crossover defect in *mlh1Δ* and the *mlh3* linker mutants, or conversely, MI nondisjunction may be more frequent than should be expected from the crossover defect in *mlh3Δ* and the *mlh1* N-terminal mutants. What could account for such behaviors? DNA joint molecules (dHJ and/or other recombination intermediates) persist longer than normal in the absence of the obligate MutLγ-accessory factor Exo1 [63]. Perhaps in *mlh1Δ* and the *mlh3* linker mutants these joint molecules persist just long enough to facilitate chromosome biorientation at metaphase I, but then at anaphase I are resolved as a mix of crossovers and noncrossovers (by Yen1, Mus81/Mms4, or Slx1/Slx4) [17, 27] and/or dissolved to form noncrossovers by Sgs1/Top3/Rmi1 [26, 29, 72]. In *mlh3Δ* and the *mlh1* N-terminal mutants in contrast, the joint molecules might be taken apart too early to facilitate chromosome biorientation, or might persist too long and thereby interfere with chromosome separation at anaphase. The possibility that alternative chromosome segregation mechanisms are at play in some MutLγ mutants, not strictly dependent on crossovers, has been noted previously based on the relatively high spore viability of an *mlh1Δ mms4Δ* strain [49]. Furthermore, the possibility that MutLγ might interfere with alternative HJ resolution systems has also been suggested previously [17].

Epistasis experiments with *mms4Δ* provide possible support for a role of joint molecules and alternative resolution pathways in explaining differences between *mlh1Δ* and *mlh3Δ* mutants. In the absence of MutLγ, Mus81/Mms4 is thought to substitute in resolving most of the joint molecules [17, 22, 23]. We speculate that Mlh1 (in the absence of Mlh3) can partially inhibit action of Mus81/Mms4, whereas Mlh3 (in the absence of Mlh1) cannot. In this model, action of Mus81/Mms4 should be more readily detected in an *mlh1Δ* mutant (where Mus81/Mms4 is not inhibited) than in an *mlh3Δ* mutant. Our observation that *mlh3Δ* is epistatic to *mms4Δ* for MI nondisjunction is thus consistent with this hypothesis. We further note that the N-terminal mutants *mlh1-K286E/R289E* and *mlh3-K316E/K320E/R323E* mutants behaved similarly to *mlh3Δ,* suggesting that these mutants may also prevent the resolution of recombination intermediates by Mus81/Mms4.

We further found that *mlh1Δ* (but not *mlh3Δ*) partially alleviated the phenotype of *msh5Δ*, in terms of both crossovers and MI nondisjunction. A similar effect was seen but not remarked on in a prior study [23]. This result can also be interpreted in terms of competition between MLH complexes and alternative resolution systems. Similar to the reasoning above, presence of MutLα and/or MutL_β_ might interfere with access of alternative crossover-promoting activities in the absence of Msh5. Absence of all MLH complexes in the *mlh1Δ* mutant is thus envisioned to allow Mus81/Mms4 and/or other activities to form more crossovers.

Taken together, these observations draw a more complex picture of the role of MutLγ in meiotic recombination, and highlight the dynamic interplay between enzymatic activities that can act on recombination intermediates.

## Materials and Methods

### Preparation of expression plasmids and baculoviruses

Oligonucleotides (oligos) used in this study were purchased from Integrated DNA Technologies. The sequence of the oligos is listed in **S5 Table**. The plasmids used in this study are listed in **S6 Table**. Vectors for expression of untagged *S. cerevisiae* Mlh1 and Mlh3 proteins were generated by cloning the corresponding gene in pFastBac1 (Invitrogen). Vectors for expression of N-terminally 6xHistidine and 2xFlag-tagged (HisFlag) Mlh1 and Mlh3 were generated by cloning the corresponding gene in pFastBac-HTbFlag, which has a sequence coding for two Flag epitopes inserted into the BamHI site of pFastBac-HTb (a gift from V. Bermudez). The *MLH1* sequence was amplified from an S288c strain (SKY4158) using primers cb111 and cb112. The PCR product was digested with BamHI and SalI restriction endonucleases and cloned into the BamHI and SalI sites of pFastBac1 to create pCCB313, or pFastBac-HTbFlag to create pCCB317. Similarly, the *MLH3* sequence was amplified from strain SKY4158 using primers cb113 and cb114. The PCR product was digested with BamHI and EcoRI and cloned into the BamHI and EcoRI sites of pFastBac1 to create pCCB318 or pFastBac-HTbFlag to create pCCB314. The cloned PCR products were verified by Sanger sequencing. The plasmid constructs coding for mutant proteins were generated by QuickChange mutagenesis using the wild-type plasmids as templates. The primers were the same ones used to create the corresponding mutant yeast strains as listed in section: Yeast strains. The viruses were produced by a Bac-to-Bac Baculovirus Expression System (Invitrogen) following the manufacturer’s instructions.

### Expression and purification of recombinant proteins

Expression of ^HisFlag^Mlh1-Mlh3 heterodimer used viruses produced from vectors pCCB313 and pCCB318. Mlh1-^HisFlag^Mlh3 heterodimer used viruses produced from vectors pCCB314 and pCCB317. *Spodoptera frugiperda* Sf9 cells were co-infected with both viruses at a multiplicity of infection of 2.5 each. Cells were harvested 62 hours after infection, washed with phosphate buffer saline (PBS), frozen in dry ice and kept at −80°C until use. A typical purification was performed with cell pellets from 500 ml culture. All the purification steps were carried out at 0– 4°C. Cell pellets were resuspended in 4 volumes of lysis buffer (25 mM HEPES-NaOH pH 7.5, 500 mM NaCl, 0.1 mM DTT, 25 mM imidazole, 1× Complete protease inhibitor tablet (Roche) and 0.1 mM phenylmethanesulfonyl fluoride (PMSF)). Cells were lysed by sonication and centrifuged at 43,000 g for 30 min. The cleared extract was re-sonicated and filtered through 0.2 μm filters before being loaded on a pre-equilibrated 1 ml HisTrap column (GE Healthcare) using an AKTA purifier (GE Healthcare). The column was washed extensively with wash buffer (25 mM HEPES-NaOH pH 7.5, 500 mM NaCl, 10% glycerol, 0.1 mM DTT, 25 mM imidazole, 0.1 mM PMSF). The tagged MutLγ complexes were then eluted with a 25–500 mM gradient of imidazole. Fractions containing MutLγ were pooled and diluted in 3 volumes of Flag buffer (25 mM HEPES-NaOH pH 7.5, 500 mM NaCl, 10% glycerol, 1 mM DTT, 2 mM EDTA). Next, the complexes were bound to 1 ml pre-equilibrated Anti-Flag M2 affinity resin (Sigma) in a poly-prep chromatography column (Bio-Rad). The resin was washed extensively with Flag buffer and the bound proteins were eluted with Flag Buffer containing 250 μg/ml 3xFlag peptide (Sigma). Fractions containing MutLγ were pooled and 200–300 μl each were loaded on 5 ml 15–40% glycerol gradients in 25 mM HEPES-NaOH pH 7.5, 400 mM NaCl, 1 mM DTT, 2 mM EDTA, 0.01% NP40. Gradients were centrifuged at 250,000 g for 20 hours and separated in about 30 fractions of 7 drops each (~175 μl). Fractions containing MutLγ were then dialyzed in 25 mM HEPES-NaOH pH 7.5, 300 mM NaCl, 10% glycerol, 1 mM DTT, 2 mM EDTA using 50 kDa cut-off Slide-a-lyzer cassettes (Thermo Scientific) and concentrated in 50 kDa cut-off Amicon centrifugal filters (Millipore). Aliquots were frozen in dry ice and stored at −80°C. Mutant MutLγ complexes were prepared using the same procedure as the wild-type proteins.

### Substrates for DNA-binding assays

DNA substrates were generated by annealing complementary oligos (sequences listed in **S6 Table**). Oligos over 40 nt were first purified on 10% polyacrylamide-UREA gels. They were subsequently mixed in equimolar concentrations (typically 10 μM) in STE (100 mM NaCl, 10 mM Tris-HCl pH 8, 1 mM EDTA), heated and slowly cooled on a PCR thermocycler (98°C for 3 min, 75°C for 1 h, 65°C for 1 h, 37°C for 30 min, 25°C for 10 min).

Substrates for electrophoretic mobility shift assays (EMSA) were assembled with the following primers: Holliday Junction (HJ^80^): cb095, cb096, cb097 and cb098; single-strand DNA (ssDNA^80^): cb099; double-strand DNA (dsDNA^80^): cb095 and cb100. Substrates were 5′ end-labeled with [γ-^32^P]-ATP (Perkin Elmer) and T4 polynucleotide kinase (New England Biolabs) and labeled substrates were purified by native polyacrylamide gel electrophoresis.

Substrates for DNA pulldown assays used the same combination of oligos as EMSA substrates except that cb095 and cb099 were replaced by 3’-biotinylated versions, cb256 and cb316, respectively. After annealing and gel purification, the substrates were bound to M280 streptavidin-coated Dynabeads (Invitrogen) in TENT buffer (10 mM Tris-HCl pH 7.5, 1 mM EDTA, 1 M NaCl and 0.1% Triton X-100) for 4 hours at 4°C. Beads were washed in TENT buffer and were stored in binding buffer (25 mM Tris-HCl pH 7.5, 10% glycerol, 100 mM NaCl, 200 μg/ml BSA, 5 mM EDTA, 2 mM DTT, 0.1% Triton X-100). Final concentration was estimated at 500 pmol DNA/μl beads (final substrate concentration of 500 nM).

Phosphorotioate-containing DNA substrates were identical to the DNA pulldown assays except that the appropriate oligos were replaced by modified versions that carried a phosphorothioate modification at the positions indicated with an asterisk in **S6 Table**. The substrate used in the FeBABE assays were (with oligos between brackets): ssDNA^-^ (cb316); ssDNA^+^ (cb361); HJ^-^ (cb256, cb96, cb97, cb98); HJ^+^ (cb256, cb351, cb97, cb98); HJ^+1^ (cb256, cb351, cb369, cb98); HJ^+2^ (cb256, cb349, cb367, cb98); HJ^+3^ (cb256, cb347, cb365, cb98); HJ^+4^ (cb256, cb345, cb363, cb98). Oligos were annealed in 40 μl STE at a concentration of 2.5 μM biotinylated oligos and a 1.2× excess of non-biotinylated oligos. Substrates were purified on a 5% Tris-acetate-EDTA-polyacrylamide gel, eluted in STE, ethanol precipitated, and resuspended in 20 mM MOPS pH 7.9.

FeBABE conjugation to phosphorothioate-containing DNA substrates were in 20 μl reactions in 20 mM MOPS pH 7.9 containing 4 μM DNA and 3.5 mM FeBABE (Dojindo). After 16 hours incubation at 50°C, the substrates were immobilized to 800 ng M280 streptavidin-coated Dynabeads (Invitrogen) in 400 μl 20 mM MOPS (pH 7.9) for 4 hours at 4°C. Excess FeBABE was removed by washing three times with 500 μl 20 mM MOPS pH 7.9 and one time with 200 μl storage buffer (25 mM Tris-HCl pH 7.5, 10% glycerol, 100 mM NaCl, 200 μg/ml BSA). Substrates were resuspended in storage buffer at an estimated concentration 1000 pmol/μl beads (final substrate concentration of about 1 μM) and stored at 4°C.

### Electrophoretic mobility shift assay

Binding reactions (20 μl) were carried out in 25 mM Tris-HCl pH 7.5, 15% glycerol, 100 mM NaCl, 2 mM DTT, 5 mM EDTA and 1 mg/ml BSA with 1 nM DNA substrate and the indicated concentrations of MutLγ. Complexes were assembled for 30 minutes at 30°C and separated on a 5% Tris-acetate-EDTA-polyacrylamide/bis (80/1) gel. Gels were dried and radioactivity was detected by phosphorimaging (Fuji).

### Protein pull-down assay

Binding reactions (20 μl) were carried out in 25 mM Tris-HCl pH 7.5, 10% glycerol, 100 mM NaCl, 2 mM DTT, 5 mM EDTA, 200 μg/ml BSA, 0.1% Triton X-100 with 25 nM immobilized DNA and the indicated concentrations of MutLγ. Complexes were assembled for 1 hour at 4°C on a rotating wheel. The beads were washed three times with 200 μl binding buffer without BSA, resuspended in 1× Laemmli sample buffer, boiled for 5 minutes and loaded on 4–12% Bis-Tris NuPAGE gels in MOPS running buffer. Proteins were detected by silver staining or anti-Flag western blotting.

### Hot substrate pull-down assay

Binding reactions (20 μl) were carried out in 25 mM Tris-HCl pH 7.5, 10% glycerol, 100 mM NaCl, 2 mM DTT, 5 mM MgCl_2_, 1 mg/ml BSA, with 25 nM biotin-labeled DNA, 25 nM ^32^P-labeled DNA and 50 nM MutLγ. Complexes were assembled for 30 minutes at 4°C. Reactions were then supplemented with 1 μl of M280 streptavidin-coated dynabeads (10 mg/ml) preequilibrated in binding buffer and 500 nM competitor dsDNA (80 bp substrate). Nucleoprotein complexes were pulled down for 1 hour at 4°C on a rotating wheel, washed three times with 200 μl of binding buffer with 0.01% NP-40 without BSA. Beads were then resuspended in loading buffer containing 0.5 mg/ml proteinase K. After 30 minutes at 30°C samples were separated on a 5% TAE-polyacrylamide gel, dried, and developed by autoradiography.

### Hydroxyl radical cleavage assay

Binding reactions (20 μl) were carried out in 25 mM Tris-HCl pH 7.5, 10% glycerol, 100 mM NaCl, 1 mg/ml BSA, 5 mM MgCl_2_ with 100 nM immobilized substrates and 100 nM Mlh1-Mlh3. Complexes were assembled for 10 minutes at room temperature, washed twice with 200 μl FeBABE buffer (25 mM Tris-HCl pH 7.5, 10% glycerol, 100 mM NaCl, 5 mM EDTA, 0.01% NP-40) with 2 mM DTT and once with 200 μl buffer without DTT. Reactions were resuspended in 20 μl FeBABE buffer and separated in two. One half (10 μl) was treated with 1.25 μl of 50 mM sodium ascorbate followed rapidly with 1.25 μl of 50 mM H_2_O_2_,10mM EDTA and the cleavage reaction performed at 30°C for 5 minutes (for ssDNA substrates) or 10 minutes (for HJ substrates). The other half was left untreated as a negative control. Reactions were quenched with 6 μl 4× LDS sample buffer and 1 μl 1 M DTT, boiled for 5 minutes and loaded on 4–12% NuPAGE Bis-Tris gels in MOPS running buffer. After electrophoresis, proteins were transferred onto Immobilon-FL PVDF membranes (Millipore) and membranes were blotted with anti-Flag M2 antibody (Sigma) followed by a IRDye 680RD goat anti-mouse IgG (Li-COR). Western blots were revealed using the Li-COR Bioscience Odyssey infrared imaging system.

### Mapping of protein-DNA interface

The molecular sizes of Flag-tagged fragments of Mlh1 and Mlh3 generated by the hydroxyl radical cleavage assay were determined by comparing their migration with MagicMark™ XP Western protein standards (Invitrogen). Profiles of each lane were quantified using ImageGauge. The peaks were identified for each molecular weight standard and used to deduce a fourth order polynomial equation of the migration distance as a function of molecular weight. The equation was used to deduce the molecular weight of the protein fragments, which in turn provide an estimate position of the hydroxyl radical cleavage sites. Based on multiple experiments we estimate that the cleavage sites are mapped within 5–10 residues.

### ATPase assay

For the Michaelis-Menten kinetics experiment, ATP hydrolysis reactions were performed with 200 nM MutLγ complexes in 25 mM Tris-HCl pH 7.5, 5 mM MgCl2, 1 mg/ml BSA and 33 nM [α^32^-P]-ATP and the indicated amounts of cold ATP. Reactions were incubated at 30°C for 30 minutes and stopped with a final concentration of 50 mM EDTA. ATP hydrolysis was measured by thin layer chromatography. Briefly, 1 μl of the reactions were spotted on a polyethyleneimine-cellulose TLC plate (EMD Chemicals) and developed in 0.5 M LiCl, 1 M formic acid. Plates were exposed to a phosphorimager screen and [α-32P]-ADP levels quantified in ImageGauge. ATPase assays in the presence of DNA had 280 nM MutLγ with 2 μM ssDNA, 1 μM dsDNA and 5 μM HJ substrates, which correspond to equal amounts (52 ng/μl) of total DNA.

### DNA cleavage assay

For nuclease assays, 20 μl reactions contained 100 nM MutLγ in the presence of 5.7 nM supercoiled substrate (200 ng of pUC19 plasmid) in 25 mM Tris-HCl pH 7.5, 5 mM MnCl_2_, 20 mM NaCl and 100 µg/μl BSA. After 1-hour incubation at 30°C, the reactions were stopped with 30 mM EDTA, 0.1% SDS and 10% glycerol. Reactions were deproteinized by treatment with 0.5 mg/ml of proteinase K for 15 min at 37°C and the DNA was separated on a 0.8% TBE-agarose gel. The gel was stained with 0.3 μg/ml ethidium bromide and imaged using a ChemiDoc MP imaging system (Bio-Rad).

### Partial Proteolysis

Trypsin digestions were performed in 30 μl reactions with 2 μM MutLγ in 25 mM HEPES-NaOH pH 7.5, 15 mM CaCl_2_, 100 mM NaCl, 3% glycerol, with or without 5 mM ATP (sodium salt, pH 7) with 25 ng trypsin (Worthington). After 5 minutes digestion at room temperature, reactions were stopped in Laemmli sample buffer. Proteins were separated on 10% NuPAGE Bis: Tris gels in MOPS SDS running buffer and visualized by Coomassie staining.

### Structural models

The structure of the C-terminal domain of Mlh1 is from PDB accession number 4e4w [73]. Homology-based models for the N-terminal domain of Mlh1 and the N and C-terminal domains Mlh3 were generated by Phyre2 [74]. The model for the MutLγ heterodimer was generated using Pymol. The PDB file is available upon request.

### AFM imaging

AFM images were captured on an Asylum Research MFP-3D-BIO (Oxford Instruments) microscope in tapping mode at room temperature. For studies of protein alone, MutLγ was diluted to a final concentration of 10 nM in 25 mM HEPES-NaOH pH 7.5, 5 mM MgCl_2_, 100 mM NaCl, 10% glycerol with or without 1 mM ATP at room temperature. A volume of 20 μl of the diluted protein was deposited onto freshly cleaved mica for 10 seconds. Samples were rinsed with 10 ml of ultrapure H_2_O and the surface was dried using a stream of nitrogen. A Bruker FMV-A AFM probe with resonance frequencies of approximately 75 kHz and spring constant of approximately 2.5 N/m was used for imaging. Images were collected at a speed of 0.8 Hz with an image size of 1 μm at 512 × 512 pixel resolution. Volume analyses of MutLγ particles were performed using a custom-written code in Metamorph. Particles with a volume between 170 and 350 nm^3^ were classified as dimers and scored as extended, one-arm folded, semi-condensed or condensed dimers based on their shape.

For protein-DNA interaction studies, reactions contained 2 nM ssDNA (~10,400 nt circle) or plasmid DNA (pUC19) with 10 or 40 nM MutLγ, respectively, in 25 mM HEPES-NaOH pH 6.8, 5 mM MgCl_2_, 50 mM NaCl and 10% glycerol. A volume of 40 μl of the reaction was deposited onto freshly cleaved mica (SPI) for 2 minutes. The sample was rinsed with 10 ml of ultrapure H_2_O and the surface was dried using a stream of nitrogen. An Olympus AC240TS-R3 AFM probe with resonance frequencies probe with resonance frequencies of approximately 70 kHz and spring constant of approximately 1.7 N/m was used. Images were collected at a speed of 0.6 Hz with an image size of 2 μm at 512 × 512 pixel resolution.

### Yeast strains and targeting vectors

Yeast strains were from the SK1 background. All the strains used in this study are listed in **S4 Table**. *mlh1Δ* and *mlh3Δ* strains were constructed by replacing the coding sequence by the *KanMX4* cassette, which was amplified from plasmid pFA6a-kanMX4 using primers cb412 and cb413 for *mlh1Δ* and cb414 and cb415 for *mlh3Δ*.

The vector used to generate the *mlh1* and *mlh3* DNA-binding mutant strains was constructed as follows. The *hphMX4* cassette was PCR amplified from pSK742 using primers cb416 and cb417 for *MLH1* and primers cb428 and cb429 for *MLH3*. The PCR products, which target *MLH1* and *MLH3* about 50 bp downstream of the corresponding gene, were transformed into fluorescent reporter strains SKY3576 and SKY3579 to achieve SKY5087 and SKY5088 for *MLH1-hphMX4* and SKY5089 and SKY5090 for *MLH3-hphMX4*. The genomic regions of *MLH1* and *MLH3* together with the downstream *hphMX4* cassettes were PCR amplified using primers cb424 and cb425 (for *MLH1*) and cb426 and cb427 (for *MLH3*) and cloned into a Topo vector using the zero Blunt Topo cloning kit (Invitrogen), creating plasmids pCCB419 (*MLH1*) and pCCB420 (*MLH3*).

Mutant plasmids were generated by QuickChange mutagenesis. Plasmid number and mutagenesis primers for each mutant are as follows: *mlh1-R214E* (plasmid pCCB423, primers cb430 and cb431); *mlh1-K253E/K254E* (plasmid pCCB424, primers cb432 and cb433); *mlh1 R273E/R274E* (plasmid pCCB425, primers cb434 and cb435); *mlh1-K286E/R289E* (plasmid pCCB426, primers cb436 and cb437); *mlh1-R341E/K344E* (plasmid pCCB427, cb438 and cb439); *mlh1-R367E/R369E/K370E/R373E* (plasmid pCCB428, primers cb470 and cb471); *mlh1-K393E/R394E* (plasmid pCCB429, primers cb442 and cb443); *mlh1-K398E/R401E* (plasmid pCCB430, primers cb444 and cb445); *mlh3-R171E/R172E/R173E* (plasmid pCCB433, primers cb454 and cb455); *mlh3-R220E/K222E* (plasmid pCCB434, primers cb456 and cb457); *mlh3-K316E/K320E/R323E* (plasmid pCCB435, primers cb458 and cb459); *mlh3-K347E/K351E* (plasmid pCCB436, primers cb460 and cb461); *mlh3-R401E/K406E/R407E* (plasmid pCCB437, primers cb462 and cb463); *mlh3-K414E/K416E* (plasmid pCCB438, primers cb464 and cb465); *mlh3-R419E/K426E* (plasmid pCCB439, primers cb466 and cb467); *mlh3-K443E/K445E/R448E* (plasmid pCCB440, primers cb468 and cb469). All mutant constructs were verified by sequencing.

Mutant strains were constructed by transformation of fluorescent reporter strains SKY3576 and SKY3579 or mismatch repair strain SKY5139 with a NheI and EcoRV (for *MLH1*) or PvuII and BglI (for *MLH3*) restriction fragment of the mutant plasmids. Selection for hygromycin resistant clones gave strains with the endogenous locus replaced with the mutant locus together with a downstream *hphMX4* cassette. Integrations at the *MLH1* locus were confirmed by Southern blotting of ApaI and SacI digested genomic DNA using a probe amplified with primers cb450 and cb451. Integrations at the *MLH3* locus were confirmed by Southern blotting of SalI and BamHI digested genomic DNA using a probe amplified with primers cb452 and cb453. The presence of the mutations was also confirmed by restriction digestion or sequencing of a PCR product amplified with primers cb111 and cb112 for *mlh1* mutants or cb113 and cb114 for *mlh3* mutants.

### Spore-autonomous fluorescence assay

All yeast cultures and sporulation were carried out at 30°C. The spore-autonomous fluorescence assay was performed as described previously [57]. Briefly, wild-type or mutant haploid strains carrying the fluorescence reporter cassettes were mated and streaked on YPD (1% yeast extract, 2% peptone, 2% dextrose, 2% agar) to isolate diploid colonies. At least three independent diploids were grown in liquid YPD overnight, transferred into YPA (1% yeast extract, 2% peptone, 2% potassium acetate) for 13.5–14 hours and sporulated in 2% potassium acetate for two days. Tetrads were then scored by fluorescence microscopy for crossovers in two test intervals and for MI-nondisjunction events. Genetic distances (cM) were calculated using the Perkins equation: cM = (100 (6NPD + TT))/(2(PD + NPD + TT)), where PD is the number of parental ditypes, NPD is the number of nonparental ditypes and TT is the number of tetratypes. Standard errors of genetic distances were calculated using the Stahl laboratory online tools (http://molbio.uoregon.edu/~fstahl/).

### Mismatch repair assay

The Lys+ reversion assay used the *lys2∷insE-A14* mutation (from strain EAY1062 from E. Alani) and the threonine reversion assay used the *hom3-10* mutation (from strain HTY1213 provided by E. Alani) [59, 61]. Mutant strains were streaked on YPD. For each mutant, at least three colonies were grown in YPD until saturation and dilutions were plated on lysine or threonine dropout medium, as appropriate, and on YPD to measure the frequencies of Lys+ or Thr+ revertants per colony-forming unit.

## Acknowledgments

We thank Eric Alani (Cornell University) for MMR reporter strains and sharing information prior to publication; Vladimir Bermudez (MSKCC) for plasmids; Navid Paknejad, Matt Brendel and Yevgeniy Romin of the Molecular Cytology Core Facility at MSKCC for assistance with AFM experiments and data analysis; and members of the Keeney lab for advice, particularly Neeman Mohibullah for suggesting the FeBABE experiment. This work received partial support from a Rapid Response Pilot Grant from the MSKCC Functional Genomics Initiative. MSKCC core facilities are supported by the NIH/NCI Cancer Center Support Grant P30 CA008748. S.K. is an Investigator of the Howard Hughes Medical Institute.

## List of Supporting information

**S1 Fig:**
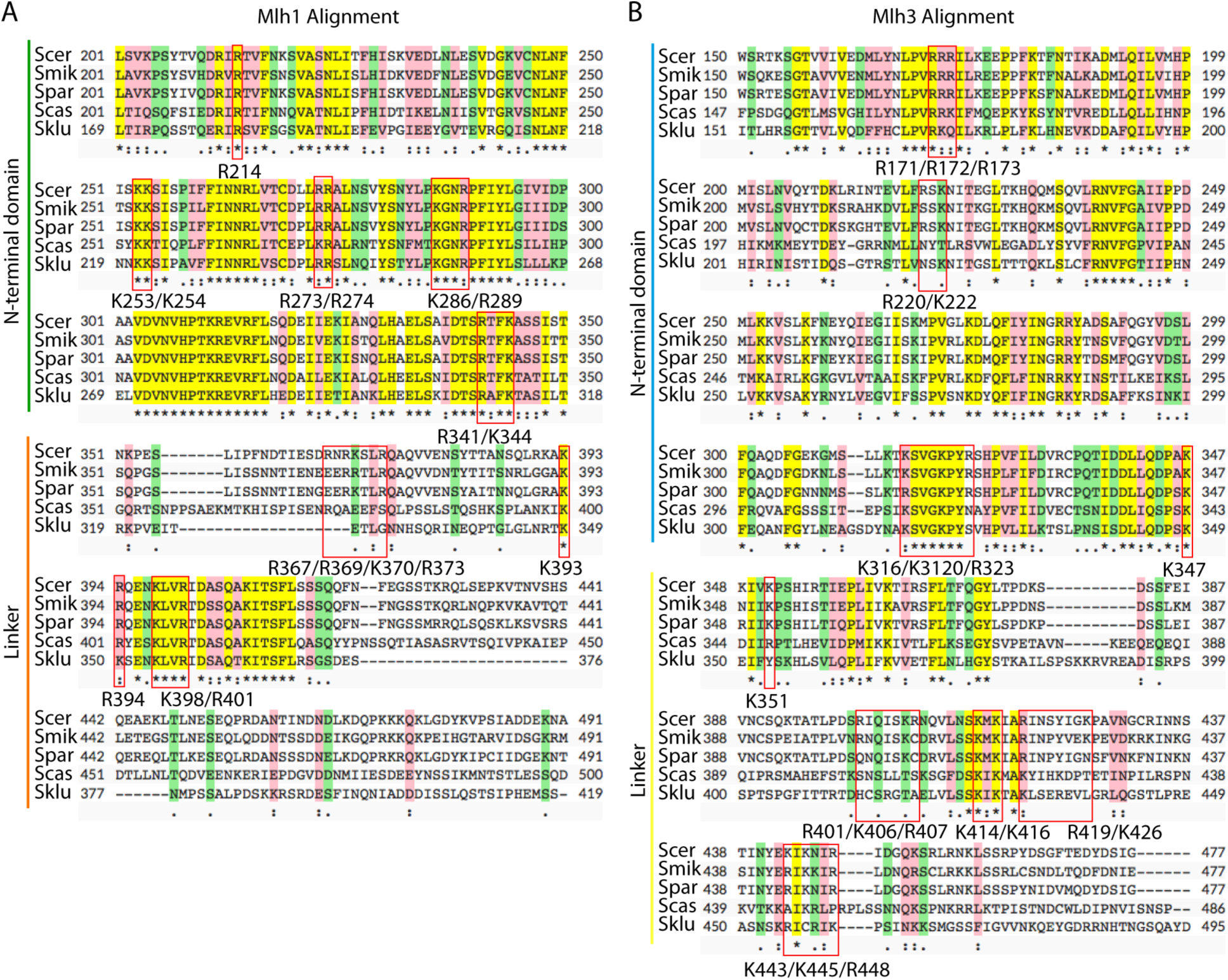
**Alignments of Mlh1 and Mlh3 from Saccharomyces species and basic residues selected for mutagenesis.** Alignments were generated using the Saccharomyces Genome Databank. Scer: *S. cerevisiae*; Smik, *S. mikatae*; Spar, *S.paradoxus*; Scas, *S. castellii*; Sklu, *S. kluyveri*. Yellow indicates conserved residues; pink indicates strong similarity; green indicates weak similarity. Red boxes highlight regions of Mlh1 (A) and Mlh3 (B) that contain lysine and arginine residues that were mutated form functional analyses.

**S1 Table:** Fluorescent spore assay data for *mlh1* and *mlh3* DNA-binding mutants

| Genotype | Strains | N | Genetic distance *CEN8-ARG4* |  |  |  | Genetic distance *ARG4-THR1* |  |  |  | MI Nondisjunctions |  |  |
| --- | --- | --- | --- | --- | --- | --- | --- | --- | --- | --- | --- | --- | --- |
|  |  |  | cM ± 95% CI | % WT ± 95% CI | p vs. WT | p vs. *mlh1Δ* | cM ± 95% CI | % WT ± 95% CI | p vs. WT | p vs. *mlh1Δ* | % ± 95% CI | p vs. WT | p vs. *mlh1Δ* |
| WT | SKY3576x3579 | 1268 | 11.63 ± 0.63 | 100.0 ± 5.42 |  |  | 4.2 ± 0.39 | 100.0 ± 9.2 |  |  | 0.08 ± 0.16 |  |  |
| *mlh1Δ* | SKY5083x5084 | 1476 | 7.22 ± 0.53 | 62.0 ± 4.56 | 3×10<sup>-9</sup> |  | 2.8 ± 0.30 | 66.6 ± 10.7 | 0.015 |  | 1.15 ± 0.54 | 4×10<sup>-4</sup> |  |
| *MLH1:Hph* | SKY5087x5088 | 666 | 10.21 ± 0.78 | 87.8 ± 6.71 | 0.32 |  | 4.6 ± 0.56 | 108.5 ± 12.2 | 0.87 |  | 0.15 ± 0.29 | 1 |  |
| *mlh1-R214E* | SKY5101x5102 | 679 | 9.50 ± 0.85 | 81.7 ± 7.31 | 0.048 |  | 3.7 ± 0.50 | 87.3 ± 13.6 | 0.71 |  | 0.00 ± 0.0 | 1 |  |
| *mlh1-K253E/K254E* | SKY5103x5104 | 687 | 4.73 ± 0.79 | 40.7 ± 6.79 | 7×10<sup>-18</sup> | 2×10<sup>-4</sup> | 2.8 ± 0.44 | 65.5 ± 15.9 | 0.057 | 1 | 2.33 ± 1.13 | <1×10<sup>-4</sup> | 0.057 |
| *mlh1-R273E/R274E* | SKY5105x5106 | 697 | 4.95 ± 0.57 | 42.6 ± 4.90 | 1×10<sup>-12</sup> | 0.02 | 1.9 ± 0.36 | 44.2 ± 19.3 | 2×10<sup>-4</sup> | 0.1 | 2.87 ± 1.24 | <1×10<sup>-4</sup> | 6×10<sup>-3</sup> |
| *mlh1-K286E/R289E* | SKY5107x5108 | 687 | 5.82 ± 0.73 | 50.1 ± 6.28 | 1×10<sup>-10</sup> | 0.17 | 2.8 ± 0.44 | 65.5 ± 15.9 | 0.057 | 1 | 3.20 ± 1.32 | <1×10<sup>-4</sup> | 1.5×10<sup>-3</sup> |
| *mlh1-R341E/K344E* | SKY5109x5110 | 675 | 6.44 ± 0.76 | 55.4 ± 6.53 | 2×10<sup>-8</sup> | 0.58 | 2.2 ± 0.40 | 52.7 ± 18.0 | 3×10<sup>-3</sup> | 0.51 | 2.22 ± 1.11 | <1×10<sup>-4</sup> | 0.082 |
| *mlh1-R367E/R369E/K370E/R373E* | SKY5111x5112 | 682 | 11.14 ± 0.89 | 95.8 ± 7.65 | 0.7 |  | 3.5 ± 0.49 | 83.4 ± 13.9 | 0.55 |  | 0.29 ± 0.4 | 0.28 |  |
| *mlh1-K393E/R394E* | SKY5113x5114 | 679 | 10.24 ± 0.77 | 88.0 ± 6.62 | 0.32 |  | 3.5 ± 0.49 | 83.8 ± 13.9 | 0.56 |  | 0.44 ± 0.5 | 0.13 |  |
| *mlh1-K398E/R401E* | SKY5115x5116 | 676 | 11.91 ± 0.82 | 102.4 ± 7.05 | 0.58 |  | 3.3 ± 0.48 | 78.9 ± 14.4 | 0.37 |  | 0.00 ± 0 | 1 |  |
| WT | SKY3576x3579 | 1268 | 11.63 ± 0.63 | 100.0 ± 5.42 |  |  | 4.2 ± 0.39 | 100.0 ± 9.2 |  |  | 0.08 ± 0.16 |  |  |
| *mlh3Δ* | SKY5085x5086 | 990 | 7.12 ± 0.62 | 61.2 ± 5.33 | 2×10<sup>-7</sup> |  | 2.6 ± 0.35 | 62.2 ± 13.3 | 0.012 |  | 2.73 ± 1.02 | <1×10<sup>-4</sup> |  |
| *MLH3:Hph* | SKY5089x5090 | 685 | 11.82 ± 0.81 | 101.7 ± 6.96 | 0.59 |  | 4.3 ± 0.54 | 102.1 ± 12.5 | 0.99 |  | 0.15 ± 0.29 | 1 |  |
| *mlh3-R171E/R172E/R173E* | SKY5121x5122 | 682 | 7.62 ± 0.69 | 65.5 ± 5.93 | 2×10<sup>-4</sup> |  | 3.6 ± 0.49 | 85.1 ± 13.6 | 0.62 |  | 1.17 ± 0.81 | 1.4×10<sup>-3</sup> |  |
| *mlh3-R220E/K222E* | SKY5123x5124 | 681 | 9.10 ± 0.74 | 78.3 ± 6.36 | 0.037 |  | 4.1 ± 0.53 | 97.4 ± 12.9 | 0.99 |  | 0.29 ± 0.4 | 0.28 |  |
| *mlh3-R316E/R320E/R323E* | SKY5125x5126 | 685 | 5.62 ± 0.72 | 48.3 ± 6.19 | 2×10<sup>-11</sup> |  | 3.1 ± 0.46 | 72.7 ± 15.0 | 0.18 |  | 3.21 ± 1.32 | <1×10<sup>-4</sup> |  |
| *mlh3-K347E/K351E* | SKY5127x5128 | 684 | 8.04 ± 0.81 | 69.1 ± 6.96 | 2×10<sup>-4</sup> |  | 3.4 ± 0.48 | 79.7 ± 14.3 | 0.40 |  | 0.88 ± 0.7 | 9×10<sup>-3</sup> |  |
| *mlh3-R401E/K406E/R407E* | SKY5129x5130 | 679 | 6.85 ± 0.77 | 58.9 ± 6.62 | 3×10<sup>-7</sup> |  | 2.5 ± 0.42 | 59.3 ± 16.8 | 0.017 |  | 1.18 ± 0.81 | 1.3×10<sup>-3</sup> |  |
| *mlh3-K414E/K416E* | SKY5131x5132 | 688 | 6.40 ± 0.64 | 55.0 ± 5.50 | 2×10<sup>-7</sup> |  | 1.7 ± 0.34 | 39.6 ± 20.3 | 3×10<sup>-5</sup> |  | 1.74 ± 0.98 | <1×10<sup>-4</sup> |  |
| *mlh3-R419E/K426E* | SKY5133x5134 | 678 | 11.28 ± 0.80 | 97.0 ± 6.88 | 0.65 |  | 4.2 ± 0.53 | 99.6 ± 12.6 | 1 |  | 0.15 ± 0.29 | 1 |  |
| *mlh3-K443E/K445E/R448E* | SKY5135x5136 | 694 | 6.99 ± 0.66 | 60.1 ± 5.67 | 6×10<sup>-6</sup> |  | 2.2 ± 0.39 | 52.9 ± 17.5 | 3×10<sup>-3</sup> |  | 0.58 ± 0.56 | 0.056 |  |
| Standard error calculation for genetic intervals was done using the Stahl lab online tool:http://molbio.uoregon.edu/~fstahl/compare2.php |  |  |  |  |  |  |  |  |  |  |  |  |  |
| p value for genetic distance were calculated by G test |  |  |  |  |  |  |  |  |  |  |  |  |  |
| p value for MI nondisjunction were calculated by Fisher's exact test (two-tailed p value) |  |  |  |  |  |  |  |  |  |  |  |  |  |

**S2 Table:** MMR data for *mlh1*and *mlh3*DNA-­binding mutants.

| Lys+ reversion | Strain | N | Frequency $\times 10^{-3}$<br>(Mean ± SD) | Frequency (normalized to<br>*mlh1*Δ) (Mean ± SD) | p |
| --- | --- | --- | --- | --- | --- |
| WT | SKY5139 | 3 | 0.1 ± 0.0 | 0.00 ± 0.001 | |
| *mlh1*Δ | SKY5175 | 6 | 43.8 ± 14.1 | 1.00 ± 0.323 | 0.0013 |
| *MLH1:Hph* | SKY5088 | 3 | 0.1 ± 0.0 | 0.00 ± 0.001 | 0.8957 |
| *mlh1-R214E* | SKY5089 | 4 | 0.5 ± 0.2 | 0.01 ± 0.004 | 0.0126 |
| *mlh1-K253E/K254E* | SKY5090 | 5 | 36.0 ± 15.5 | 0.82 ± 0.354 | 0.0081 |
| *mlh1-R273E/R274E* | SKY5091 | 6 | 36.0 ± 9.5 | 0.82 ± 0.217 | 0.0004 |
| *mlh1-K286E/R289E* | SKY5092 | 6 | 29.4 ± 12.0 | 0.67 ± 0.275 | 0.0047 |
| *mlh1-R341E/K344E* | SKY5093 | 3 | 31.0 ± 1.9 | 0.71 ± 0.045 | 0.0001 |
| *mlh1-R367E/R369E/K370E/R373E* | SKY5094 | 5 | 0.6 ± 0.1 | 0.01 ± 0.002 | 0.0002 |
| *mlh1-K393E/R394E* | SKY5095 | 6 | 21.1 ± 9.9 | 0.48 ± 0.226 | 0.0092 |
| *mlh1-K398E/R401E* | SKY5096 | 6 | 32.0 ± 17.2 | 0.73 ± 0.392 | 0.0172 |
| Lys+ reversion | Strain | N | Frequency $\times 10^{-6}$<br>(Mean ± SD) | Frequency (normalized to<br>*mlh3*Δ) (Mean ± SD) | p |
| WT | SKY5139 | 6 | 33 ± 3 | 0.19 ± 0.045 | |
| *mlh3*Δ | SKY5176 | 6 | 172 ± 22 | 1.00 ± 0.320 | <0.0001 |
| *MLH3:Hph* | SKY5199 | 6 | 42 ± 5 | 0.24 ± 0.068 | 0.0036 |
| *mlh3-R171E/R172E/R173E* | SKY5200 | 6 | 161 ± 22 | 0.94 ± 0.318 | <0.0001 |
| *mlh3-R220E/K222E* | SKY5201 | 6 | 151 ± 4 | 0.88 ± 0.063 | <0.0001 |
| *mlh3-R316E/R320E/R323E* | SKY5202 | 6 | 182 ± 15 | 1.06 ± 0.215 | <0.0001 |
| *mlh3-K347E/K351E* | SKY5203 | 5 | 131 ± 9 | 0.76 ± 0.113 | <0.0001 |
| *mlh3-R401E/K406E/R407E* | SKY5204 | 6 | 98 ± 4 | 0.57 ± 0.064 | <0.0001 |
| *mlh3-K414E/K416E* | SKY5205 | 6 | 89 ± 4 | 0.52 ± 0.062 | <0.0001 |
| *mlh3-R419E/K426E* | SKY5206 | 6 | 89 ± 6 | 0.52 ± 0.088 | <0.0001 |
| *mlh3-K443E/K445E/R448E* | SKY5207 | 6 | 61 ± 7 | 0.36 ± 0.104 | <0.0001 |
| Thr+ reversion | Strain | N | Frequency $\times 10^{-6}$<br>(Mean ± SD) | Frequency (normalized to<br>*mlh1*Δ) (Mean ± SD) | p |
| WT | SKY5137 | 3 | 0 ± 0 | 0.00 ± 0.000 | |
| *mlh1*Δ | SKY5173 | 3 | 102 ± 7 | 1.00 ± 0.067 | <0.0001 |
| *MLH1:Hph* | SKY5177 | 3 | 0 ± 0 | 0.00 ± 0.000 | |
| *mlh1-R214E* | SKY5178 | 3 | 1 ± 1 | 0.01 ± 0.006 | 0.1583 |
| *mlh1-K253E/K254E* | SKY5179 | 3 | 53 ± 9 | 0.52 ± 0.090 | 0.0005 |
| *mlh1-R273E/R274E* | SKY5180 | 3 | 88 ± 18 | 0.86 ± 0.173 | 0.0011 |
| *mlh1-K286E/R289E* | SKY5181 | 3 | 46 ± 6 | 0.46 ± 0.063 | 0.0002 |
| *mlh1-R341E/K344E* | SKY5182 | 3 | 65 ± 12 | 0.64 ± 0.114 | 0.0007 |
| *mlh1-R367E/R369E/K370E/R373E* | SKY5183 | 3 | 0 ± 0 | 0.00 ± 0.003 | |
| *mlh1-K393E/R394E* | SKY5184 | 3 | 35 ± 2 | 0.35 ± 0.022 | <0.0001 |
| *mlh1-K398E/R401E* | SKY5185 | 3 | 76 ± 3 | 0.74 ± 0.029 | <0.0001 |
| N is the number of independent cultures.<br>p values were calculated by unpaired t test. |  |  |  |  |  |

**S3 Table:** Fluorescent spore assay data in *mms4Δ* and *msh5Δ* mutant backgrounds.

| Genotype | Strains | N | Genetic distance *CEN8-ARG4* |  | MI Nondisjunctions<br>% ± 95% CI |
| --- | --- | --- | --- | --- | --- |
|  |  |  | cM ± 95% CI | % WT ± 95% CI |  |
| WT | SKY3576x3579 | 1268 | 11.63 ± 0.63 | 100.0 ± 5.4 | 0.08 ± 0.16 |
| *mms4Δ* | SKY5531x5532 | 689 | 11.47 ± 0.89 | 98.6 ± 7.7 | 0.87 ± 0.69 |
| *mlh1Δ* | SKY5083x5084 | 1476 | 7.22 ± 0.53 | 62.0 ± 4.6 | 1.15 ± 0.54 |
| *mlh1Δ mms4Δ* | SKY5558x5559 | 595 | 3.19 ± 0.50 | 27.5 ± 4.3 | 3.03 ± 1.38 |
| *mlh3Δ* | SKY5085x5086 | 990 | 7.12 ± 0.62 | 61.2 ± 5.3 | 2.73 ± 1.02 |
| *mlh3Δ mms4Δ* | SKY5564x5565 | 601 | 2.50 ± 0.64 | 21.5 ± 5.5 | 3.00 ± 1.36 |
| *mlh1-Nterm* | SKY5107x5108 | 687 | 5.82 ± 0.73 | 50.1 ± 6.3 | 3.20 ± 1.32 |
| *mlh1-Nterm mms4Δ* | SKY5560x5561 | 522 | 1.92 ± 0.42 | 16.5 ± 3.6 | 4.02 ± 1.69 |
| *mlh3-Nterm* | SKY5125x5126 | 685 | 5.62 ± 0.72 | 48.3 ± 6.2 | 3.21 ± 1.32 |
| *mlh3-Nterm mms4Δ* | SKY5566x5567 | 601 | 1.25 ± 0.32 | 10.7 ± 2.8 | 3.33 ± 1.43 |
| *mlh1-linker* | SKY5113x5114 | 679 | 10.24 ± 0.77 | 88.0 ± 6.6 | 0.44 ± 0.5 |
| *mlh1-linker mms4Δ* | SKY5562x5563 | 311 | 7.07 ± 0.99 | 60.8 ± 8.5 | 2.89 ± 1.86 |
| *mlh3-linker* | SKY5131x5132 | 688 | 6.40 ± 0.64 | 55.0 ± 5.5 | 1.74 ± 0.98 |
| *mlh3-linker mms4Δ* | SKY5568x5569 | 600 | 1.33 ± 0.33 | 11.5 ± 2.8 | 2.83 ± 1.33 |
| WT | SKY3576x3579 | 1268 | 11.63 ± 0.63 | 100.0 ± 5.4 | 0.08 ± 0.16 |
| *msh5Δ* | SKY3580x3581 | 685 | 3.50 ± 0.75 | 30.1 ± 6.4 | 13.72 ± 2.58 |
| *mlh1Δ* | SKY5083x5084 | 1476 | 7.22 ± 0.53 | 62.0 ± .6 | 1.15 ± 0.54 |
| *mlh1Δ msh5Δ* | SKY5570x5571 | 600 | 4.50 ± 0.58 | 38.7 ± 5.0 | 8.67 ± 2.25 |
| *mlh3Δ* | SKY5085x5086 | 990 | 7.12 ± 0.62 | 61.2 ± 5.3 | 2.73 ± 1.02 |
| *mlh3Δ msh5Δ* | SKY5576x5577 | 602 | 3.16 ± 0.81 | 27.1 ± 7.0 | 13.62 ± 2.74 |
| *mlh1-Nterm* | SKY5107x5108 | 687 | 5.82 ± 0.73 | 50.1 ± 6.3 | 3.20 ± 1.32 |
| *mlh1-Nterm msh5Δ* | SKY5572x5573 | 601 | 3.33 ± 0.68 | 28.6 ± 5.8 | 11.81 ± 2.58 |
| *mlh3-Nterm* | SKY5125x5126 | 685 | 5.62 ± 0.72 | 48.3 ± 6.2 | 3.21 ± 1.32 |
| *mlh3-Nterm msh5Δ* | SKY5578x5579 | 593 | 2.61 ± 0.46 | 22.5 ± 4.0 | 11.64 ± 2.58 |
| *mlh1-linker* | SKY5113x5114 | 679 | 10.24 ± 0.77 | 88.0 ± 6.6 | 0.44 ± 0.5 |
| *mlh1-linker msh5Δ* | SKY5574x5575 | 600 | 2.92 ± 0.48 | 25.1 ± 4.1 | 9.83 ± 2.38 |
| *mlh3-linker* | SKY5131x5132 | 688 | 6.40 ± 0.64 | 55.0 ± 5.5 | 1.74 ± 0.98 |
| *mlh3-linker msh5Δ* | SKY5580x5581 | 600 | 2.00 ± 0.40 | 17.2 ± 3.4 | 13.67 ± 2.75 |
| N is the number of tetrads scored. |  |  |  |  |  |

**S4 Table:**
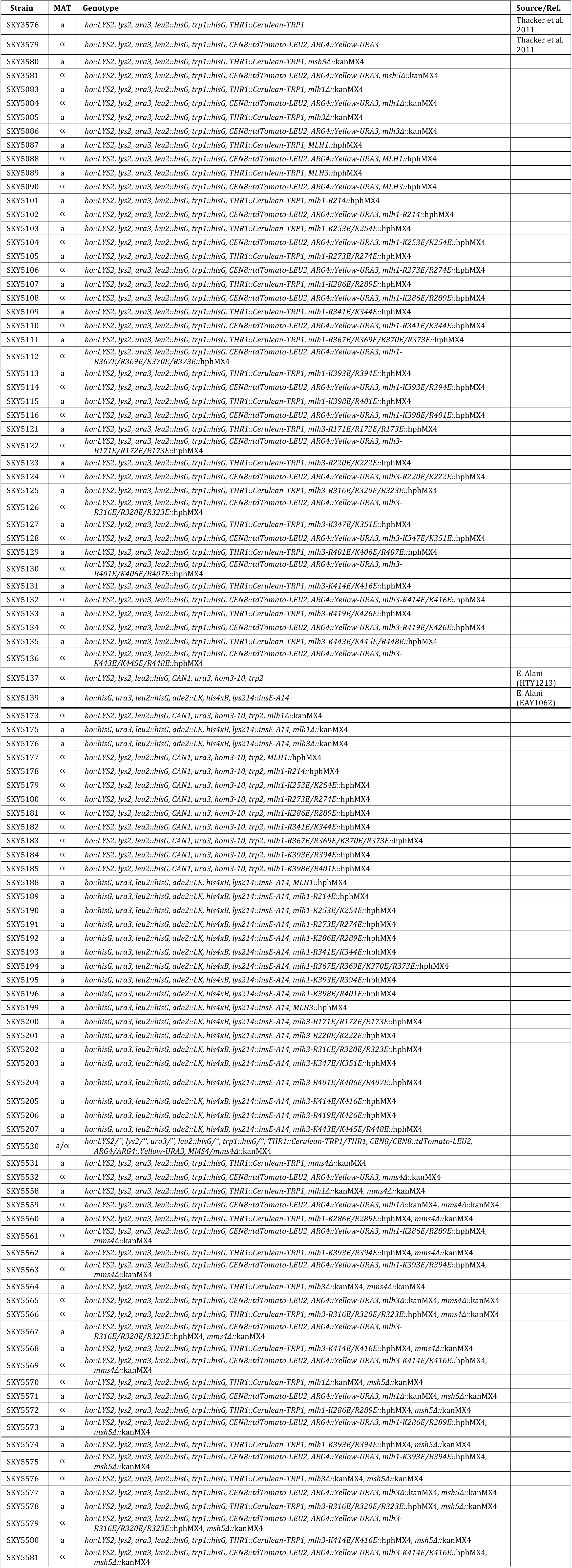
List of yeast strains.

**S5 Table:**
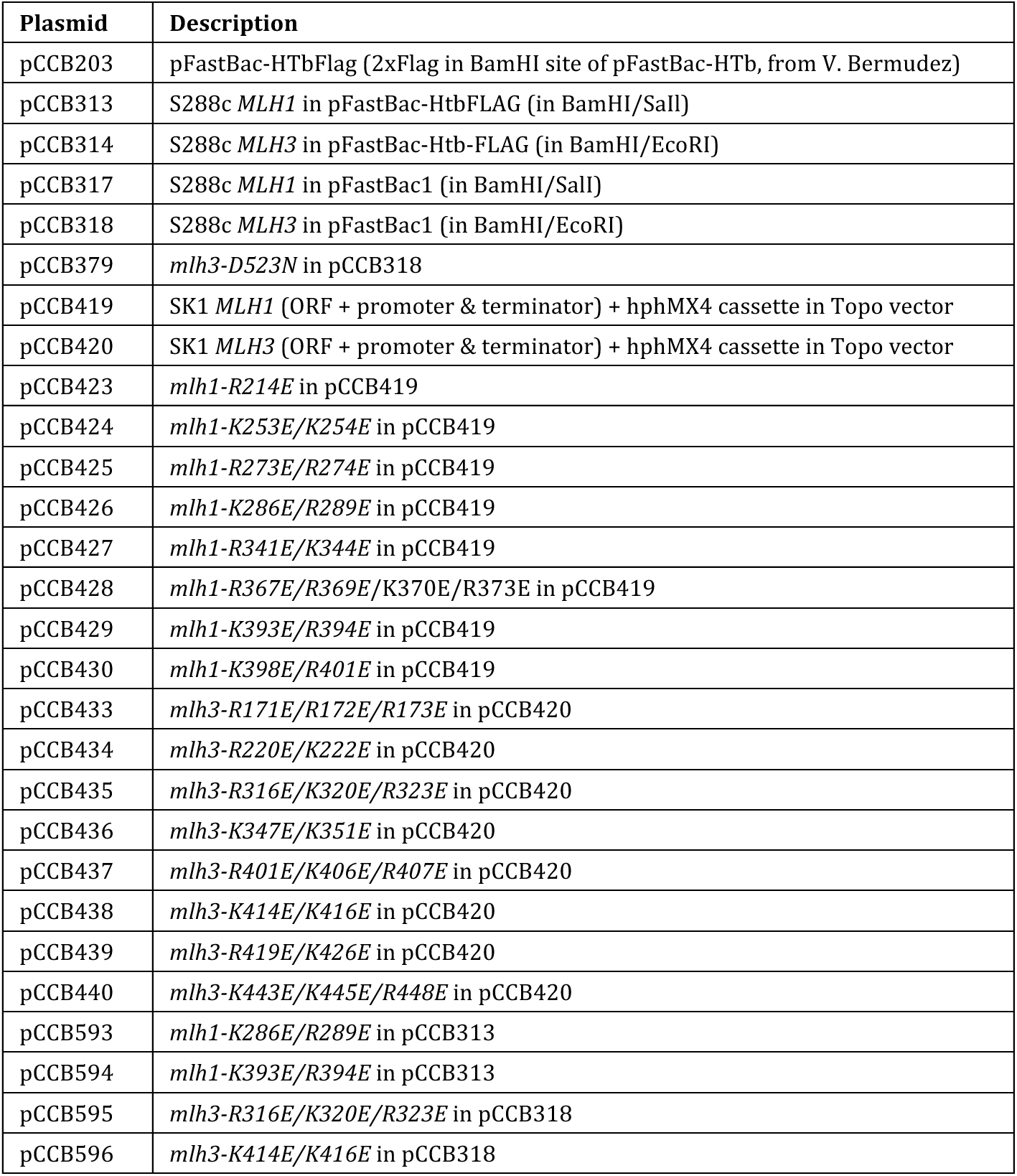
List of plasmids.

**S6 Table:**
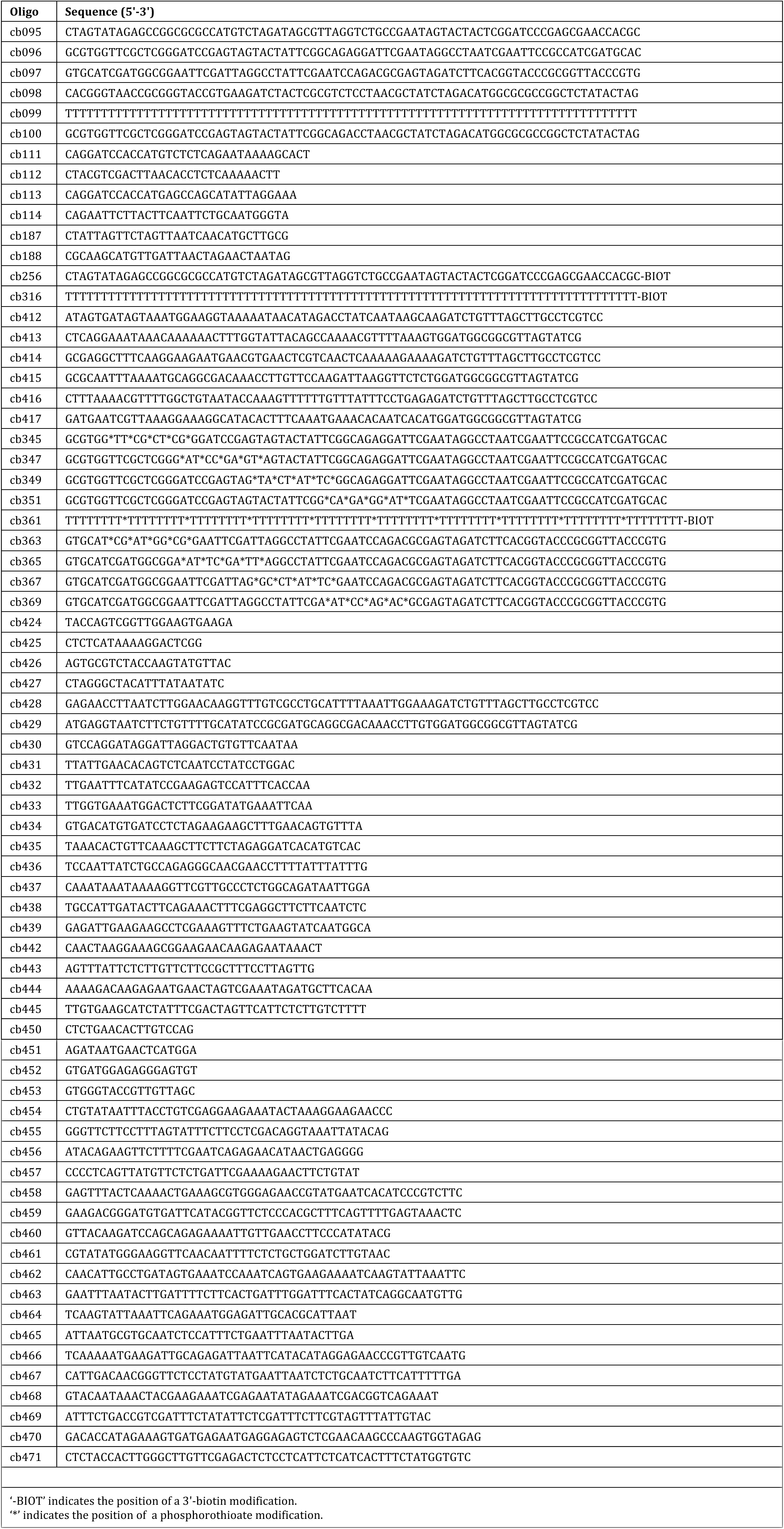
List of oligonucleotides.

